# The serine protease homolog Skanda modulates Toll-Phenoloxidase-mediated immunity in *Drosophila*

**DOI:** 10.1101/2025.09.30.679548

**Authors:** Sanjana Vasanth, Yang Wang, Nathan Klotz, Anzer Khan, Chao Xiong, Jean-Philippe Boquete, Tisheng Shan, Prince Kumar Sah, Haobo Jiang, Bruno Lemaitre

**Author notes:** Corresponding author Bruno Lemaitre.

## Abstract

Extracellular serine protease (SP) cascades are central regulators of insect innate immunity. These cascades are negatively controlled by serine protease inhibitors (serpins) and fine-tuned by serine protease homologs (SPHs), which resemble SPs but lack catalytic activity. In *Drosophila*, a key SP cascade—the Toll–phenoloxidase (PO) pathway—governs both the melanization response and Toll-dependent antimicrobial peptide production. This cascade is triggered by secreted pattern-recognition receptors or microbial proteases and converges on two clip-domain SPs, Persephone and Hayan, which activate the Toll ligand Spätzle via the Spätzle-Processing Enzyme (SPE) and process prophenoloxidases. Here, we characterize the SPH Skanda and uncover its role in the Toll–PO cascade. *skanda* is genomically clustered with *hayan* and *persephone* and is transcriptionally induced upon infection. Skanda is unusual among SPHs, containing a long serine/threonine-rich region, two clip domains, and atypical disulfide bonds. Skanda is unstable and subject to cleavage by Grass. Functional assays show that Skanda dampens activation of Hayan, and to a lesser extent Persephone, within the Toll–PO cascade. Notably, *skanda*-deficient flies are highly susceptible to *Staphylococcus aureus* despite displaying normal Toll signaling and cuticular melanization. Moreover, compound mutants lacking two members of the *hayan–psh–skanda* cluster reveal a hidden contribution of Skanda to Toll activation in the absence of Persephone. Together, our results identify Skanda as a modulatory SPH that fine-tunes Toll pathway activity in concert with Persephone and Hayan.

## Introduction

Extracellular serine protease (SP) cascades play a crucial role in regulating insect innate immune responses, (Jiang and Kanost, 2000; Veillard et al., 2016; Westlake et al., 2024; Zhao et al., 2011). These cascades rely on a core mechanism in which multiple SPs are sequentially activated through zymogen cleavage. This stepwise activation connects pathogen recognition to effector responses, shaping the overall immune reaction. In *Drosophila*, a common SP cascade, the Toll-phenoloxidase (Toll-PO) SP cascade regulates two immune processes: the melanization reaction and Toll pathway-mediated production of antimicrobial peptides (Westlake et al., 2024).

Melanization is an arthropod-specific immune response that involves the deposition of melanin at wound sites. It contributes to wound healing and prevents the entry of microbes into the hemocoel (Nakhleh et al., 2017). The reactive oxygen species (ROS) produced during the process are directly toxic to microbes (Tang, 2009; Westlake et al., 2024; Zhao et al., 2011, 2007). A rate-limiting enzyme in the formation of melanin is phenoloxidase (PO), which catalyzes the conversion of phenols to quinones.

These quinones then polymerize to form melanin. POs are produced in an inactive form called prophenoloxidases (PPOs). Three PPOs have been identified in *Drosophila*; PPO1 and PPO2, produced by crystal cells, are involved in hemolymph melanization, while PPO3, produced by lamellocytes, is involved in the encapsulation of parasite eggs (Binggeli et al., 2014; Dudzic et al., 2015; Nam et al., 2008). Upon injury, crystal cells burst to release PPO1 and PPO2 into the hemolymph (Bidla et al., 2007; Rizki et al., 1980). PPO1 and PPO2 are activated upon cleavage by SPs in the end of the Toll-PO SP cascade, which also triggers the Toll pathway (Dudzic et al., 2019; Shan et al., 2023; Westlake et al., 2024).

In *Drosophila*, the Toll pathway is essential for defense against Gram-positive bacteria and fungi, controlling the expression of antimicrobial peptides in fat body (Lemaitre et al., 1996; Rutschmann et al., 2002; Ryckebusch et al., 2025). Unlike mammalian Toll-like receptors (TLRs), which directly recognize microbial patterns as pattern recognition receptors (PRRs), the *Drosophila* Toll receptor is activated indirectly by a cleaved form of the secreted protein Spätzle (Spz) (Lemaitre et al., 1996; Weber et al., 2003). The SPs involved in proSpz cleavage during immune responses differ from those responsible for its processing in the dorsoventral patterning of the early embryo (Dudzic et al., 2019; Stein et al., 2013; Valanne et al., 2011; Westlake et al., 2024). Among them, the Spz Processing Enzyme (SPE), which contains a clip domain, has been identified as the final SP responsible for cleaving proSpz and activating the Toll pathway (Jang et al., 2008, 2006; Jiang and Kanost, 2000; Kambris et al., 2006). Consistent with biochemical studies done in other insects (Jiang et al., 2009; Kim et al., 2008), genetic analysis supports the presence of one core SP cascade in Drosophila, the Toll-PO SP cascade, that connects microbial recognition to Toll pathway activation on one hand and activation of melanization on the other (Dudzic et al., 2019; Shan et al., 2023; Westlake et al., 2024) (**Figure 1**). Upstream of the Toll-PO SP cascade, secreted pattern recognition receptors (PRRs) detect microbial cell wall components : GNBP1 and PGRP-SA recognize peptidoglycans from Gram-positive bacteria, while GNBP3 detects fungal β-1,3-glucan (Gobert et al., 2003; Gottar et al., 2006; Leulier et al., 2003; Michel et al., 2001; Pili-Floury et al., 2004; Westlake et al., 2025). Upon binding, these PRRs cause autoactivation of modular serine protease (ModSP), which then would activate cSP48 to mature Grass. Grass activation ultimately leads to the activation of two serine proteases Persephone (Psh) and Hayan, which redundantly regulate SPE activation leading to the maturation of the Toll ligand Spz and activating PPO1 and PPO2 leading to melanization (Buchon et al., 2009; Dudzic et al., 2019; El Chamy et al., 2008; Gobert et al., 2003; Gottar et al., 2006; Pili-Floury et al., 2004).The Toll-PO SP cascade can be directly activated at the level of Psh-Hayan, through direct cleavage of the Psh protease bait region by microbial subtilisin or proteases with the help of the cathepsin CtsK1 (Issa et al., 2018). This triggers SPE activation, leading to Toll pathway activation by Spätzle (Spz) cleavage and to melanization reaction (El Chamy et al., 2008; Issa et al., 2018; Ligoxygakis et al., 2002; Ming et al., 2014; Nakano et al., 2023; Nam et al., 2012). Most of the SP involved in the Toll-PO SP cascade contain a clip domain, which is thought to regulate the sequential activation through zymogen cleavage by upstream proteases (Jang et al., 2008; Piao et al., 2005; Veillard et al., 2016). A serine protease Sp7 has been implicated in the melanization reaction in larvae and adults, but Sp7 deficient flies still melanize around the wound (Castillejo-López and Häcker, 2005; Dudzic et al., 2019, 2015). Genetic approach has allowed to distinguish two activities associated with PPO1 and PPO2: cuticular melanization at the injury site which is independent of Sp7 and an uncharacterized microbicidal activity against *Staphylococcus aureus* which is dependent on Sp7 (Dudzic et al., 2019). Recent biochemical studies have identified additional serine proteases that await genetic characterization (Jin et al., 2023; Shan et al., 2023). Moreover, the Drosophila genome encodes many other clip-domain serine proteases, some being induced upon infection, that have not yet been characterized (Cao et Jiang 2018; De Gregorio, Spellman, et al. 2002).

**Figure 1.**
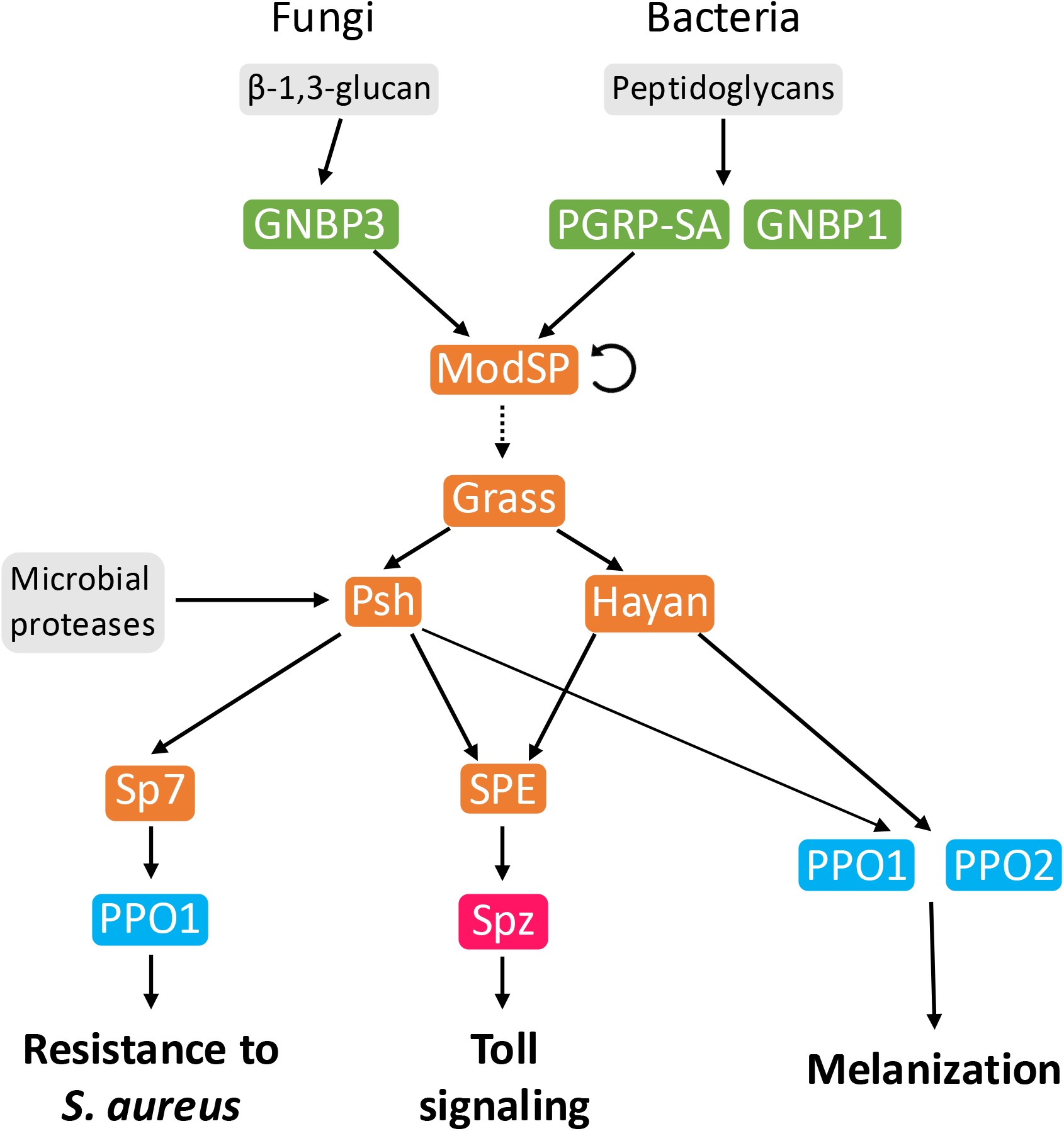
A simplified model of the Toll-PO serine protease cascade. Systemic infection of *Drosophila* activates the Toll-PO serine protease (SP) cascade in the hemolymph. Lysine-type peptidoglycans from Gram-positive bacteria and β-glucans from fungi are detected by pattern recognition receptors (PRRs) PGRP-SA, GNBP1 and GNBP3, respectively. This triggers the sequential activation of SPs ModSP, Grass, and Psh/Hayan. Psh and Hayan mediate the activation of SP SPE, which activates by cleavage the Toll ligand Spz, activating an intracellular cascade leading to the production of antimicrobial peptides (middle). Psh (and likely Hayan) can also be independently cleaved by microbial proteases. *S. aureus* host defense is mediated by the activation of SP Sp7 and prophenoloxidase-1 (PPO1 (left). Hayan and Psh also activate PPO1 and PPO2, initiating the melanization response leading to cuticular blackening (right). In green: PRRs, orange: SPs, pink: cytokines, blue: PPOs. Inspired from (Westlake et al., 2024)

Hayan and Psh are two SPs that are central to the Toll-PO cascade (Dudzic et al., 2019) (**Figure 1**). They are encoded by paralogous genes located only 751 bp apart on the *Drosophila melanogaster X* chromosome. Hayan and Psh form a closely related monophyletic lineage among other CLIP SPs. Bioinformatic evidence indicates that Psh evolved specifically in the Melanogaster group of flies from a gene duplication event of an ancestral Hayan gene that is conserved across the *Drosophila* genus. Hayan also has a protease “bait region” similar to Psh suggesting that it could also activated by microbial protease (Dudzic et al., 2019; Shan et al., 2023). Flies lacking Hayan failed to show cuticular melanization but display wild-type Toll activity indicating an essential role of this SP in activating PPO1 and PPO2 (Dudzic et al., 2019; Nakano et al., 2023; Nam et al., 2012). In contrast, flies lacking *psh* display normal activation of the Toll pathway by the pattern recognition receptor PGRP-SA and GNBP1, but have reduced Toll activity (Buchon et al., 2009; El Chamy et al., 2008; Ligoxygakis et al., 2002). The use of *Hayan, psh* double mutant has revealed that they play overlapping and redundant functions: flies lacking both *Hayan* and *psh* have strongly reduced Toll pathway activity and melanization reaction upon both pattern-recognition activation and microbial protease injection. This led to the notion that Psh-Hayan form a signaling hub that integrates signals from pathogens to activate through distinct serine proteases Toll pathway activation and melanisation (Dudzic et al., 2019).

The gene *CG15046* (hereafter *skanda*) is located in the same gene locus as *Hayan* and *psh*, only 529 bp away from *psh. skanda* encodes an uncharacterized CLIP serine protease homolog (SPH). SPHs are serine protease-like proteins mutated in their catalytic residues. They have a regulatory function (Ross et al., 2003). Like *Hayan* and *psh*, *skanda* gene expression is upregulated in *Drosophila* upon infection and regulated by the Toll pathway (De Gregorio *et al*., 2002b). The genomic localization of Skanda within the *Hayan* and *psh* gene cluster and its inducibility by the Toll pathway points to a role in *Drosophila* innate immunity. Here we have performed a biochemical and genetic functional characterization of Skanda in *Drosophila* host defense. Our study shows that Skanda modulates Toll signaling activity with Psh and Hayan. Strikingly, *skanda* deficient flies displayed a marked susceptibility to infection with the Gram-positive bacterium *S. aureus*, albeit displaying nearly wild-type Toll activation and cuticular melanization. Compound mutants deleting two of three SP of the *Hayan-psh-skanda* gene cluster reveal a cryptic role of Skanda in Toll activation in absence of Psh. Thus, we characterized the serine protease homolog Skanda as a new component of the Toll-PO SP cascade that shapes the *Drosophila* immune responses.

## Results

### Skanda is an unusual clip SPH regulated by the Toll and Imd pathways

*CG15046* is an immune induced gene regulated by the Toll and Imd pathways (E. De Gregorio et al., 2002b) (**Figure 2A**), which is clustered with Hayan and Psh on the *X* chromosome. As Hayan and Persephone are the names of Korean and Greek gods, respectively; we decided to rename CG15046 Skanda after an Indian god. *Drosophila* Skanda is a 494-residue protein with a predicted signal peptide (residues 1−25) for secretion (**Figure 2B)**. The mature protein contains a first clip domain (34−83), a Serine-Threonine (ST)-rich region (84−164), a second clip domain (167−216), a linker (217−250), and a serine Protease-Like Domain (PLD, 251−481). S1A peptidase domains are characterized by the perpendicular arrangement of two adjacent β-barrel-like structures (Page and Di Cera, 2010). Each barrel is made up of six antiparallel β-strands. The His-Asp-Ser catalytic triad responsible for SP’s proteolytic activity is located in the cleft formed between the barrels (Bode and Schwager, 1975; Huber et al., 1974; Veillard et al., 2016; Wang et al., 1985). To confirm that Skanda is an SPH, we generated a 3D model of Skanda and superimposed it to the known structure of Grass, a clip-domain SP (**Figure 2C**)(Jumper et al., 2021; Kellenberger et al., 2011; Pettersen et al., 2004). The peptidase and peptidase-like domains of the two proteins overlapped greatly, confirming that Skanda is a S1A family member. Close inspection of the catalytic triad showed that the catalytic residues of Skanda, Gly^296^-Asp^332^-Val^444^ are not conserved. The replacements of His with Gly and Ser with Val, and Asp sidechain pointing away from the Gly abolish the peptidase activity. Sequence comparison indicates that the *N*-terminal regions of Skanda’s PLD (Y^261^–G^378^ and R^253^–R^393^) are ∼40% similar to the corresponding segments of Persephone (Psh; Y^155^–G^274^) and Hayan (R^131^–R^273^), respectively. Although the remaining PLD sequences diverge from canonical protease domains, AlphaFold modeling predicts that Skanda retains a similar overall fold. Five disulfide bonds (C^281^-C^297^, C^358^-C^458^, C^385^-C^437^, C^400^-C^418^, and C^440^-C^470^) stabilize the PLD, two of which appear to be unique (**Figure 2B**, *red* lines). Unlike cSPH35 or cSPH242 (Jin et al., 2023), Skanda lacks a key Cys residue in the linker so that, if the precursor is cleaved between A^250^ and I^251^, the N-terminal fragment of Skanda is not covalently tethered to the PLD. If Skanda is cleaved by a protease at the hydrophilic interdomain regions, the protein may dissociate into 2 to 3 fragments due to their structural instability. In contrast, if Skanda is only cleaved within a clip domain, the protein remains attached by the three intradomain disulfide bonds. A conformational change induced by cleavage within a clip or protease-like domain may be needed for Skanda’s regulatory function.

**Figure 2.**
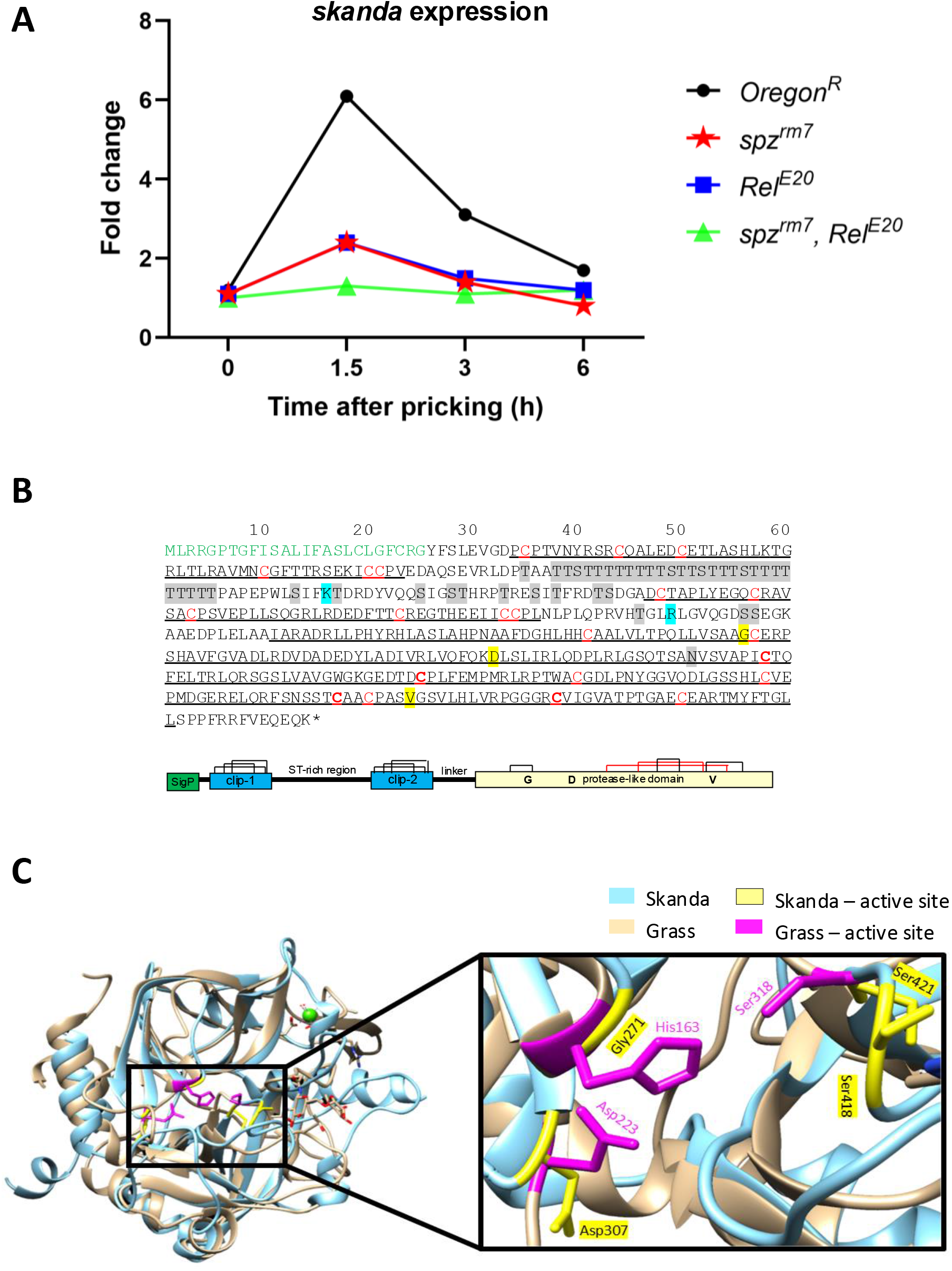
Skanda is a CLIP-SPH whose expression is regulated by the Toll and Imd pathways. **A.** *skanda* is an early immune-induced gene, with expression peaking at 1.5 hours post-pricking with a mixture of *M. luteus* and *E. coli*. *skanda* expression is regulated by the Toll and Imd pathways, as *skanda* upregulation upon infection was lost in *spz^rm7^* and *Relish^E20^*flies. OregonR was used as the wild-type control. Data from (Ennio De Gregorio et al., 2002). The data are extracted from Affymetrix array dataset from De Gregorio et al., done with polyA extracts of male flies pricked with a mixture of the Gram-positive bacterium *M. luteus* and the Gram-negative bacterium *E. coli*. **B.** Structural features of *D. melanogaster* proSkanda. Upper panel: Signal peptide in *green* font. Clip-1, clip-2, and protease-like domain sequences are underlined. Consistent with AlphaFold prediction of the 3D structure, 18 of the 22 Cys residues (in *red* font) form nine disulfide bonds, well conserved in clip-domain proteases. The remaining 4 Cys residues (in *bold*) are also predicted to form two unique disulfide bonds. Putative *N*- and *O*-linked glycosylation sites are shaded *gray*. The Gly, Asp, and Val residues, shaded *yellow*, correspond to the catalytic triad His, Asp, and Ser in an S1A serine protease. Lower panel: Signal peptide, clip domains, and protease-like domain are shown as *green*, *blue*, and *yellow* boxes, respectively. The conserved and unique Cys residues are connected with *black* and *red* lines. The ST-rich region, linker, and other structures are shown proportionally. Based on the results of mass spectrometric analysis (Fig. S1, Table S1) and SDS-PAGE analysis (Fig. 3), Skanda may be cleaved at or before Lys^125^ and Arg^218^ (in *cyan*) by Grass, cSP48, and a trypsin-like protease released by Sf9 cells. **C.** Overlap of the peptidase domain of Grass (in *brown*) and peptidase-like domain of Skanda (in *blue*) visualized using Chimera v1.17.3. The domains overlap greatly, confirming that Skanda is a S1A family member. Inset: The catalytic triads of Grass (in *magenta*) and Skanda (in *yellow*). The replacements of His with Gly and Ser with Val, and Asp sidechain pointing away from the Gly abolish the peptidase activity.

We conclude that Skanda is an unusual SPH due to the presence of a long ST domain, two clip domains, and some atypical disulfide linkages. The presence of clip domains, the clustering with Hayan and Psh, the immune inducibility, all point to a role in *Drosophila* immunity.

### Proteolytic processing of the Skanda by SP48 and Grass

To investigate Skanda’s biochemical properties, we expressed the full-length precursor with affinity tags (**Table S2**) in *Sf9* cells, an insect ovarian cell line derived from *Spodoptera frugiperda,* using a baculovirus system. We purified 0.663 mg of recombinant protein from 1 liter of culture. SDS-PAGE analysis revealed bands at ∼85, 44, and 35 kDa (**Figure 3A**), indicating partial proteolytic processing during expression and purification. The ∼30 kDa difference between the observed and theoretical molecular mass (54,599 Da) suggests significant *O*-linked glycosylation, particularly in the ST-rich region next to clip domain-1. The first cleavage event appears to occur near Arg^154^, upstream of clip domain-2, generating a 44 kDa product with a smeared appearance−larger than the calculated 36,788 Da−consistent with glycosylation. A second cleavage around Arg^223^ yields a 35 kDa doublet, exceeding the theoretical mass of 31,252 Da. These discrepancies further support the extensive *O*-linked glycosylation.

**Figure 3.**
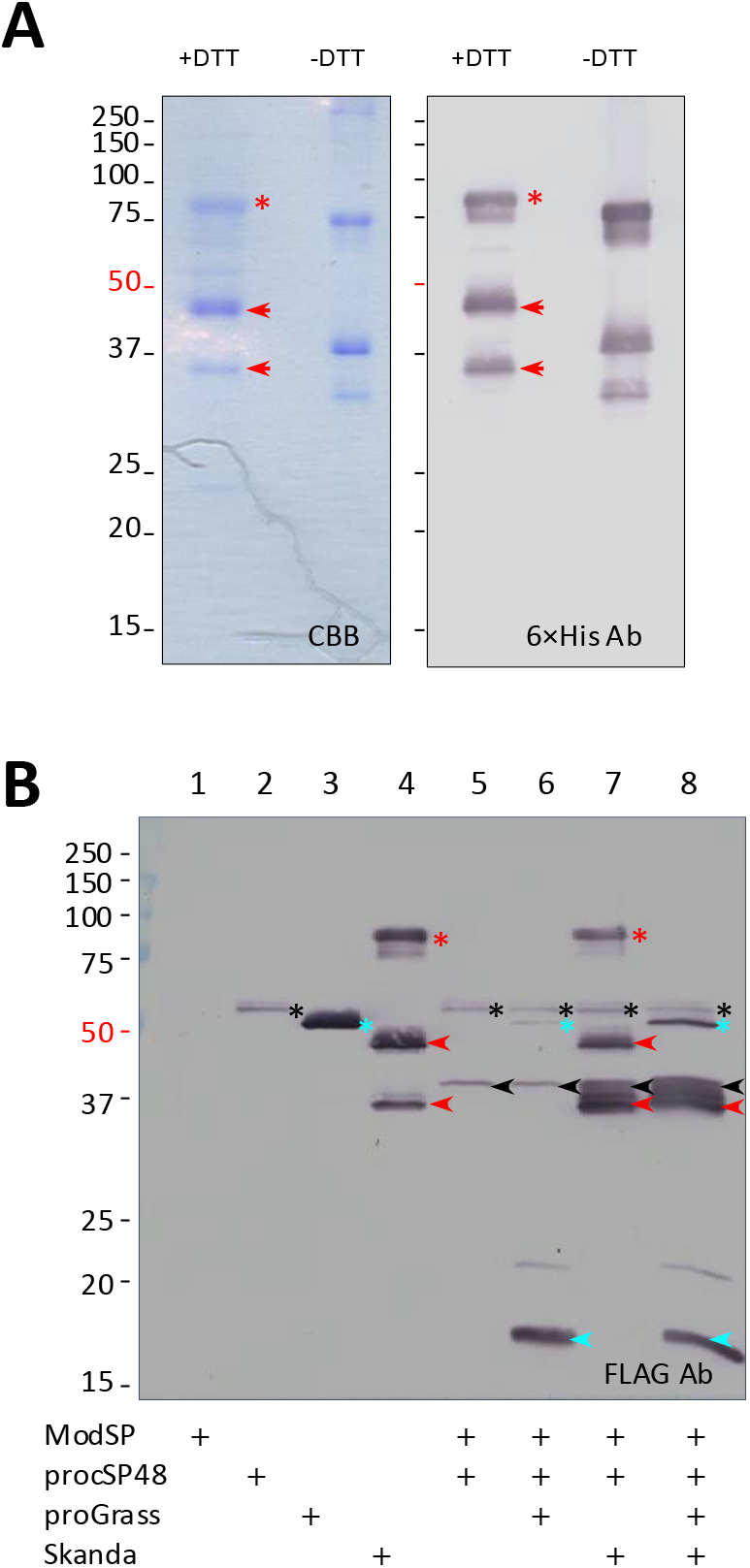
Skanda is an unstable SPH that can be processed by Grass as well as cSP48. **A.** SDS-PAGE analysis of the Skanda, a mixture of proSkanda and its fragments. The purified sample was treated with SDS sample buffer with or without dithiothreitol (DTT), separated on a 12% polyacrylamide gel, stained with Coomassie brilliant blue (CBB, 2 μg protein, *left* panel) or analyzed by immunoblotting (0.2 μg protein, *right* panel) using diluted antiserum to 6×His tag. The proSkanda and its cleavage products are marked with *red asterisk* and *arrowheads*, respectively. Positions and sizes (in kDa) of the pre-stained *M*_r_ standards are indicated on the *left*, with the 50 kDa marker highlighted *red*. **B.** Immunoblot analysis of the recombinant Skanda cleaved by *D. melanogaster* Grass. *Drosophila* ModSP (0.5 μg), proSP48 (0.1 μg), proGrass (0.4 μg), and Skanda (1 μg) were incubated in 25 μl of buffer A at 37 °C for 1 h. The reaction and controls were subjected to 12% reducing SDS-PAGE and immunoblot analysis using 1:1000 diluted antiserum against the FLAG tag, located at SP48 and Skanda C-termini and Grass N-terminus (Shan et al., 2023). ModSP has a C-terminal 6×His tag. Precursors (*) and major cleavage products (*arrowheads*) of SP48, Grass, and Skanda are in *black*, *cyan*, and *red*, respectively. Based on the mass spectrometric analysis (Fig. S1, Table S1) and SDS-PAGE analysis (Fig. 3), K125 and R218 may represent the cleavage sites of cSP48, Grass, and a trypsin-like protease released by Sf9 cells.

Under nonreducing conditions, the Skanda bands shift to lower apparent molecular weights (∼75, 40, and 31 kDa), implying that cleavage did not take place within clip domains between the conserved Cys-3 and Cys-4 residues. This contrasts with the cleavage behavior observed in cSPH35 and cSPH242 (Jin et al., 2023), where intradomain disulfide linkages retain covalently associated fragments that migrate at ≥75 kDa. Our previous study demonstrated that ModSP activates proSP48, SP48 activates proGrass, and Grass activates proPersephone and proHayan-PA/PB (Shan et al., 2023). To determine whether Grass could also cleave Skanda, which shares homology with Psh and Hayan, we incubated ModSP, proSP48, proGrass, and Skanda individually or in combination. Reactions and controls were separated by 12% SDS-PAGE, and cleavage products were analyzed using an anti-FLAG antibody (**Table S2**). ProSP48 (52 kDa, lane 2) was largely converted to active SP48, with its catalytic domain migrated at ∼38 kDa (lane 5) (**Figure 3B**). SP48 efficiently cleaved proGrass (50 kDa, lane 3) and also acted on Skanda (85 and 44 kDa), producing active Grass (lane 6 vs. lane 3) and a greater abundance of 35−38 kDa Skanda (lane 7 vs. lane 4). The N-terminal light chain of Grass (12.2 kDa) appeared as a predominant 17 kDa band recognized by FLAG antibody (lane 6). When proGrass was added to the reaction (lane 7), the 85 and 44 kDa Skanda bands were completely converted to the 35−38 kDa range (lane 8), suggesting that Grass processed proSkanda and its 44 kDa intermediate at multiple sites. The intensity of cleavage products in the 36−37.5 kDa region was notably higher in lane 8 compared to lane 7. We used LC-MS/MS analysis to further characterize the cleavage of Skanda, Although the processing events were confirmed semi-quantitatively (**Figure 3B**), we could not fully define the precise cleavage sites (**Supp. Figure S1, Supp. document S1**).

We conclude that Skanda can be processed by Grass in the Toll-PO SP cascade which is consistent with Skanda acting at the same level of Hayan and Psh in the proteolytic cascade.

### Effect of Skanda on proPsh, proHayan-PA, and proHayan-PB cleavage by Grass

To evaluate whether the presence of Skanda influences Grass-mediated processing of Psh and Hayan precursors, we incubated ModSP, proSP48, proGrass, and the precursors of Psh, Hayan-PA, or Hayan-PB with or without Skanda. A small portion of proPsh was converted into faint bands at 32, 29 and 26 kDa (**Figure 4A**, *left*, lane 1 vs. lane 4), and the addition of Skanda further reduced the intensity of these bands (lane 4 vs. lane 6). We observed that the processing of proHayan-PA and -PB by ModSP, proSP48, proGrass into their catalytic domains (indicated by *arrowheads*) was more efficient than proPsh, particularly for Hayan-PB (**Figure 4A**, *middle*). Skanda addition led to a noticeable reduction in the 36 kDa band corresponding to the catalytic domain (**Figure 4A**, *right*, lane 2 vs. lane 3; lane 5 vs. lane 6). As Skanda concentration increased from 0 to 0.2, 0.4, and 0.8 μg, the intensity of the catalytic domain bands decreased proportionally (**Figure 4A***, right*), indicative of dose-dependent inhibition. Consistent with a negative impact of Skanda on the cleavage of Hayan and Psh, Skanda tends to reduce the cleavage of downstream SPs SP7, MP1, SPE and Ser7 (**Supp. Figure S2**, **Supp. document S2**).

**Figure 4.**
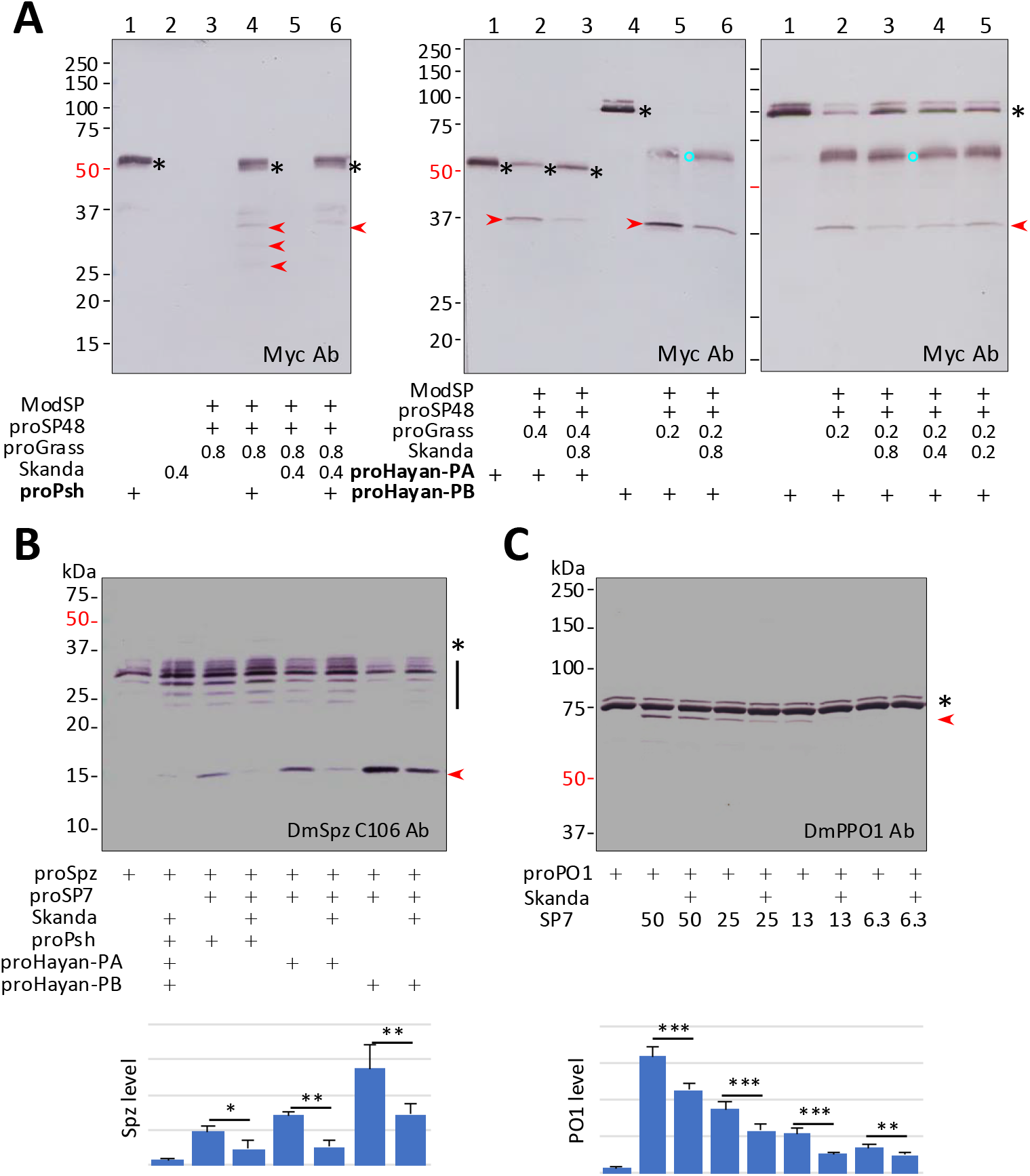
Skanda negatively regulates the processing of proPSH and Hayan by upstream serine proteases. **A.** Product analysis of proPersephone (*left*), proHayan-PA (*middle*) and -PB (*right*) cleaved by Grass in the absence and presence of Skanda. *Drosophila* ModSP (0.5 μg), proSP48 (0.1 μg), proGrass (0.8, 0.4, or 0.2 μg), Skanda (0.2. 0.4, or 0.8 μg), proPsh (0.25 μg) or proHayan-PA/PB (0.1 μg) were incubated in 25 μl buffer A at 37 °C for 1 h. The reactions and controls were subjected to 12% reducing SDS-PAGE and immunoblot analysis using 1:1000 diluted antibody against the c-Myc tag, located at C-terminus of Hayan. Precursors (*), intermediate (○), and major cleavage products (*arrowheads*) of Psh and Hayan are in *black*, *cyan*, *red*, respectively. **B.** Effect of the Skanda on the cleavage activation of *Drosophila* proSpӓtzle by SP7. ModSP (0.5 μg), proSP48 (0.1 μg), proGrass (0.1 μg), Skanda (1.0 μg), proPsh or proHayan-PA/PB (0.1 μg), proSP7 (0.4 μg), and proSpӓtzle (0.4 μg) were incubated in 25 μl buffer A at 37 °C for 1 h. The reaction and controls were subjected to 15% reducing SDS-PAGE and immunoblot analysis using 1:2000 diluted anti-Spӓtzle C106 antibody. Precursor, intermediates, and Spӓtzle are marked with *asterisk*, *vertical line*, and *arrowhead*, respectively. Relative levels of Spӓtzle or integrated intensities of the 15 kDa band on four immunoblot images are analyzed using ImageJ and plotted as mean ± SEM (n = 4). Student’s t-test (two-tailed, type-2) was used to analyze the statistical significance of level differences in pairwise comparisons. **C.** Effect of the Skanda on the cleavage activation of *Drosophila* of proPO1 by SP7. ModSP (0.5 μg), proSP48 (0.1 μg), proGrass (0.1 μg), Skanda (1.0 μg), proPsh (0.1 μg), proSP7 (50, 25, 12.5, and 6.25 ng), proPO1 (0.4 μg), were incubated in 25 μl buffer A at room temperature for 1 h. The reactions and controls were subjected to 7.5% reducing SDS-PAGE and immunoblot analysis using 1:2000 diluted antibody against *Drosophila* proPO1. Precursor (*) and 72 kDa PO1 (*red arrowhead*) are indicated. Relative levels of the 70 kDa PO1 bands on four immunoblot are analyzed as described above. *, *p* <0.05; **, *p* <0.01; ***, *p* <0.001.

### Effect of Skanda on proSpz and proPO1 cleavage activation

*Drosophila* SP7, SPE, MP1, and Ser7 are known to activate proSpätzle (proSpz), while SP7 and MP1 also activate proPO1 and proPO2 (Shan et al., 2023). To explore whether Skanda modulates these downstream immune responses, we assessed its effect on SP7-mediated activation of proSpz and proPO1 *in vitro*., The 15 kDa active Spätzle fragment was detected following incubation of ModSP with cSP48, Grass, Psh, SP7, and proSpz, as visualized using an antibody against Spätzle C106 (**Figure 4B**). Substituting proPsh with proHayan-PA or -PB led to enhanced production of the active form of Spätzle. However, Skanda addition reduced Spätzle production, consistent with decreased levels of Psh and Hayan observed in **Figure 4A**. SP7, once activated by the ModSP-cSP48-Grass module, cleaved proPO1 in a concentration-dependent manner (**Figure 4C**). The addition of Skanda reduced the intensity of the 72 kDa PO1 band, indicating decrease in proPO1 activation. Collectively our biochemical analysis reveals that Skanda tends to negatively impact the activation of Psh and Hayan and consequently the downstream SP, and ultimately the cleavage of Spz and PPO.

### *skanda* is not essential for the activation of the Toll pathway

After the biochemical study, we initiated a genetic analysis of the role of Skanda in *Drosophila* immunity. For this, we generated *skanda^Δ107^*flies carrying an 8 bp deletion causing a frameshift in the second exon using CRISPR/Cas9 (**Figure 5A**, **Supp. Figure 3A**). This mutation was then isogenized for seven generations into the *w^1118^ DrosDel* isogenic genetic background as described in (Ferreira et al., 2014). *skanda^Δ107^* flies were homozygous viable with no obvious external deformities. To rule out any cis effects caused by the mutation, we tested *Hayan* and *psh* expression in *skanda^Δ107^* flies. Flies had wild-type expression of *Hayan* and *psh*, confirming that the mutation does not affect these flanking SP genes (**Supp. Figure 3B**). We did not find any impact of Skanda mutation in the regulation of the Imd pathway **(Supp. Figure 3C and 3D, Supplementary document S3**). This result was not surprising as the Imd pathway is not known to be regulated by serine proteases, but by intracellular and membrane-bound PGRP receptors (Ferrandon et al., 2007; Westlake et al., 2024). It however reveals that loss of *skanda* does not lead to a general weakness of the fly that could impact its host defense.

**Figure 5.**
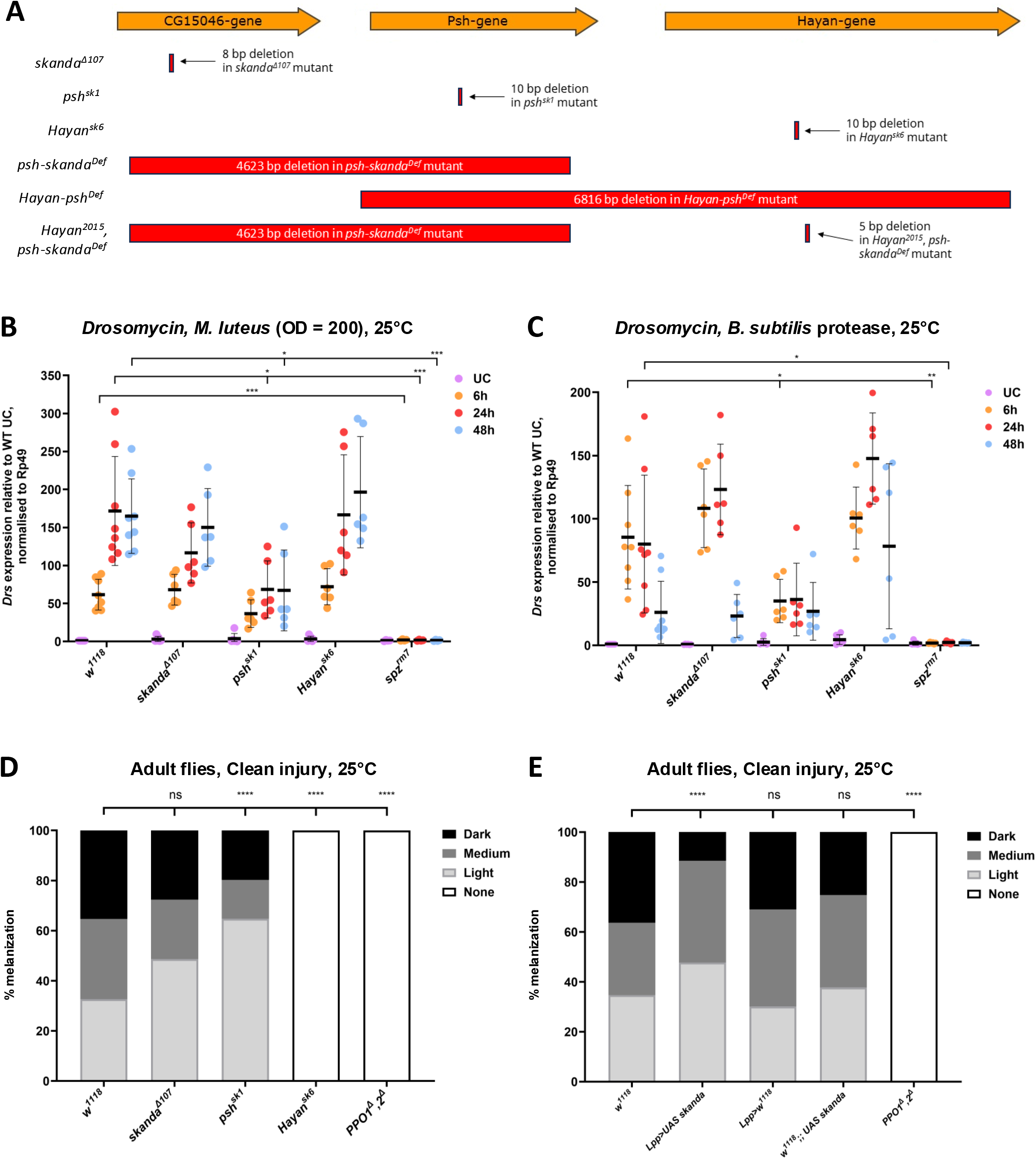
Skanda is not a critical component of the Toll and melanization cascades. (**A**) The *Hayan-psh-skanda* gene cluster is located on the *X* chromosome. *psh* is located 751 bp away from *Hayan*, and *skanda* is located 529 bp away from *psh*. Shown in red are the deletions generated in this study removing *skanda* and *psh* (*psh-skanda^Def^)* and *psh* and *Hayan* (*Hayan-psh^Def^*). The triple mutant was produced by inducing a mutation in *Hayan* in *psh-skanda^Def^*. (**B, C**). *Drs* expression of *skanda* mutants was observed 6, 24 and 48 hours after (**B**) pricking with *M. luteus* OD_600_ 200 and (**C**) injection with *B. subtilis* protease. *spz^rm7^*flies were used as a positive control. *skanda^Δ107^* and *Hayan^sk6^* flies showed wild-type induction of *Drs*, while *psh^sk1^* flies showed reduced *Drs* expression. As expected, *Drs* expression was strongly reduced in *spz^rm7^* flies. Mixed-effects analysis indicated that there was a significant difference in *Drs* expression over time across genotypes (*p* < 0.0001 for both time and genotype factors). (**D**) Cuticular melanization of *skanda* mutants was observed 16 hours post-clean injury with a sterile needle. Dark, medium and light refers to the size and color of the scab (see **Supp. Figure 5A**). *PPO1^Δ^,2^Δ^* flies were used as a positive control. *skanda^Δ107^* flies showed wild type melanization, while *psh^sk1^* flies showed reduced melanization. *Hayan^sk6^* flies showed no response to clean injury. As expected, *PPO1^Δ^,2^Δ^* flies showed no melanization. (**E**) Cuticular melanization of flies over-expressing *skanda* with the *Lpp-Gal4* driver was observed 16 hours post-clean injury with a sterile needle. Dark, medium and light refers to the size and colour of the scab (see **Supp. Figure 5A**). As expected, *PPO1^Δ^,2^Δ^*flies showed no melanization. ns = not significant (not shown for 5B, C), *p < 0.05, **p < 0.01, ***p < 0.001, and ****p < 0.0001.

We then turned our attention to the Toll pathway, which is regulated by extracellular serine proteases including Hayan and Psh. The Toll pathway can be activated by microbial cell wall components such as peptidoglycans or glucans that are recognized by PRRs, or by microbial proteases (Gottar et al., 2006; Leulier et al., 2003). It regulates the expression of a large set of antimicrobial peptides and Bomanins, including Drosomycin, an antifungal peptide which is often used as read-out to monitor this pathway (Hanson et al., 2025; Neyen et al., 2014; Troha and Buchon, 2019). We monitored the expression of *Drosomycin* in wild-type, *skanda^Δ107^*and *spz^rm7^* flies upon infection with the avirulent Gram-positive bacterium *Micrococcus luteus*, which is known to stimulate the PRR but not the Psh branch of the Toll-PO SP cascade (Buchon et al., 2009; Gottar et al., 2006). In our experiments, we also included flies carrying null mutations in *psh* and *Hayan*, affecting the two SP-coding genes clustered with *skanda*. *skanda^Δ107^* flies showed wild-type induction of *Drs*, while as expected, *Drs* expression was strongly reduced in *spz* flies (**Figure 5B**). To our surprise, we observed a reduced expression of *Drs* expression in *psh^sk1^* but not *Hayan^sk6^* flies 24 and 48 hours post-infection with *M. luteus*. This suggests that Psh is also required for Toll activation by PRR and is not fully redundant with Hayan (Dudzic et al., 2019)(**Figure 5B**). Consistent with significant Toll activity, *skanda^Δ107^* flies show only an intermediate susceptibility to the Gram-positive bacterium *Enterococcus faecalis* and the yeast *Candida albicans,* that are mostly controlled by the Toll pathway (Gottar *et al., 2006;* Rutschmann *et al., 2002;* Ryckebusch *et al., 2025*) (**Supp. Figure 3E and F)**.We then generated *UAS-Skanda* transgenic flies to see whether the over-expression of Skanda could affect Toll pathway activation by *M. luteus*. We validated the *UAS-Skanda* insertion, but noticed that the insertion was leaky resulting in a significantly higher expression of Skanda without any Gal4 (**Supp. Figure 4A**). Over-expression of *skanda* alone or using the fat body *Lpp-Gal4* driver did not affect the expression of *Drs* compared to the wild-type in *M. luteus* infected flies (**Supp. Figure 4B**).

In addition to sensing microbial cell wall components through the pattern recognition receptors PGRP-SA and GNBP3, the Toll pathway can be directly activated by microbial proteases at the level of Psh (El Chamy et al., 2008; Gottar et al., 2006; Ming et al., 2014). We therefore analyzed whether the loss or over-expression of *skanda* could impact the activation of the Toll pathway upon injection of *B. subtilis* proteases. *skanda^Δ107^* and *Hayan^sk6^* flies showed wild-type *Drs* expression, while, as expected, *psh^sk1^*flies showed significantly reduced *Drs* expression 6 hours post-injection (**Figure 5C**). Thus, our studies did not reveal any major role of *skanda* in the activation of the Toll pathway by either the protease or PRR branches.

### *skanda* is not a critical component of cuticular melanization

The Toll-PO cascade also regulates PPO activation, leading to melanin production. We therefore hypothesized that Skanda could be involved in the activation of melanization and not the Toll pathway. Thus, we investigated whether *skanda* was required for melanization of the cuticle upon injury. Adult flies were first pricked with a sterile needle of diameter 0.2 mm, a stimulus that triggers a blackening reaction at the wound site. Based on the size and color of the black spot, flies were classified as having dark, medium, light or no melanization (**Supp. Figure 5A**). *PPO1^Δ^,2^Δ^* flies, lacking two PO genes, were used as a positive control as they are incapable of cuticular melanization (Binggeli et al., 2014; Dudzic et al., 2015). *skanda^Δ107^*flies had a wild-type melanization response, while *PPO1^Δ^,2^Δ^*flies showed no melanization (**Figure 5D**). This experiment also confirms the critical role of Hayan in cuticular melanization as *Hayan^sk6^* showed almost no blackening. Interestingly, *psh^sk1^* mutants have slightly reduced blackening. The melanization reaction at the wound site is stronger in presence of microbial elicitors such as *M. luteus*. We therefore monitored cuticular melanization upon septic injury with *M. luteus*. Although the blackening reaction was stronger, we did not observe any significant difference between the wild-type and *skanda^Δ107^* flies (**Supp. Figure 5C**). Similar results were obtained at a third instar larval stage, when injury triggers a marked melanization (**Supp. Figure 5B and 5D)**). Interestingly, we however noticed that overexpressing *skanda* using a fat body *Lpp-Gal4* driver or in absence of Gal4 caused the flies to have a reduced melanization response to clean injury, but not to septic injury with *M. luteus* (**Figure 5E, Supp. Figure 5E**). Thus, consistent with the biochemical analysis that reveals an inhibition of Hayan by Skanda, we observe an inhibition of the cuticular melanization reaction by Skanda, but restricted to clean injury.

### *skanda* mutants are highly susceptible to *S. aureus* infection

We recently developed an adult infection model using a low-dose inoculation of the Gram-positive bacterium *S. aureus*, which is especially appropriate to study melanization. Strikingly, flies deficient for PPOs, but not flies lacking hemocytes or Toll effectors, rapidly succumb to this challenge (Dudzic et al., 2019; Hanson et al., 2019; Ryckebusch et al., 2025). The melanization response restricts the growth of *S. aureus*, preventing its systemic dissemination. Furthermore, Dudzic et al., revealed a disconnect between survival to infection and the blackening of the wound site (Dudzic et al., 2019). Although a mutation in *Hayan* leads to the almost complete loss of the blackening reaction in adults, *Hayan* mutants do not share the susceptibility of *PPO1,2* mutant flies against *S. aureus*. In contrast, flies mutated for another serine protease Sp7 do not survive *S. aureus* infection, despite almost wild-type levels of cuticle blackening in adult. Thus, a third mechanism of defense exists downstream of the Toll-PO cascade that likely relies on the PPOs, but does not encompass cuticular melanization or the Toll pathway (Dudzic et al., 2019; Westlake et al., 2024). This third mechanism has not yet been characterized, although the production of ROS by POs, to which *S. aureus* is very susceptible (Ramond et al., 2021), has been invoked. A previous study noted that *skanda* was induced in hemocytes upon *S. aureus* infection (Nazario-Toole, 2016). This prompted us to analyze whether *skanda* mediates immune response to *S. aureus*. Adult flies were pricked with *S. aureus* at low dose (OD_600=_ 0.1). *PPO1^Δ^,2^Δ^* flies, completely lacking melanization, were used as a positive control as they are known to rapidly succumb to this challenge. *Bom^Δ55C^* flies were used as a control for the Toll pathway response, as they are almost as susceptible as *spz^rm7^* flies when infected with Gram-positive bacteria. We observed that *skanda^Δ107^* flies were highly susceptible to *S. aureus* infection, similar to *PPO1^Δ^,2^Δ^*flies (**Figure 6A**). This was a surprising result, as we have shown above that these flies have mostly normal Toll and cuticular melanization responses. *Hayan^sk6^* and *psh^sk1^*flies displayed an intermediate susceptibility, similar to *Bom^Δ55C^*flies. Overexpressing *skanda* using the *Lpp-Gal4* fat body driver significantly increased the survival of flies compared to the wild-type. Even in absence of Gal4 driver, the significant expression of Skanda by the *UAS-Skanda* insertion provided protection (**Figure 6B, Supp. Figure 4A**). This reinforces the notion that Skanda has a specific role in resisting *S. aureus* infection.

**Figure 6.**
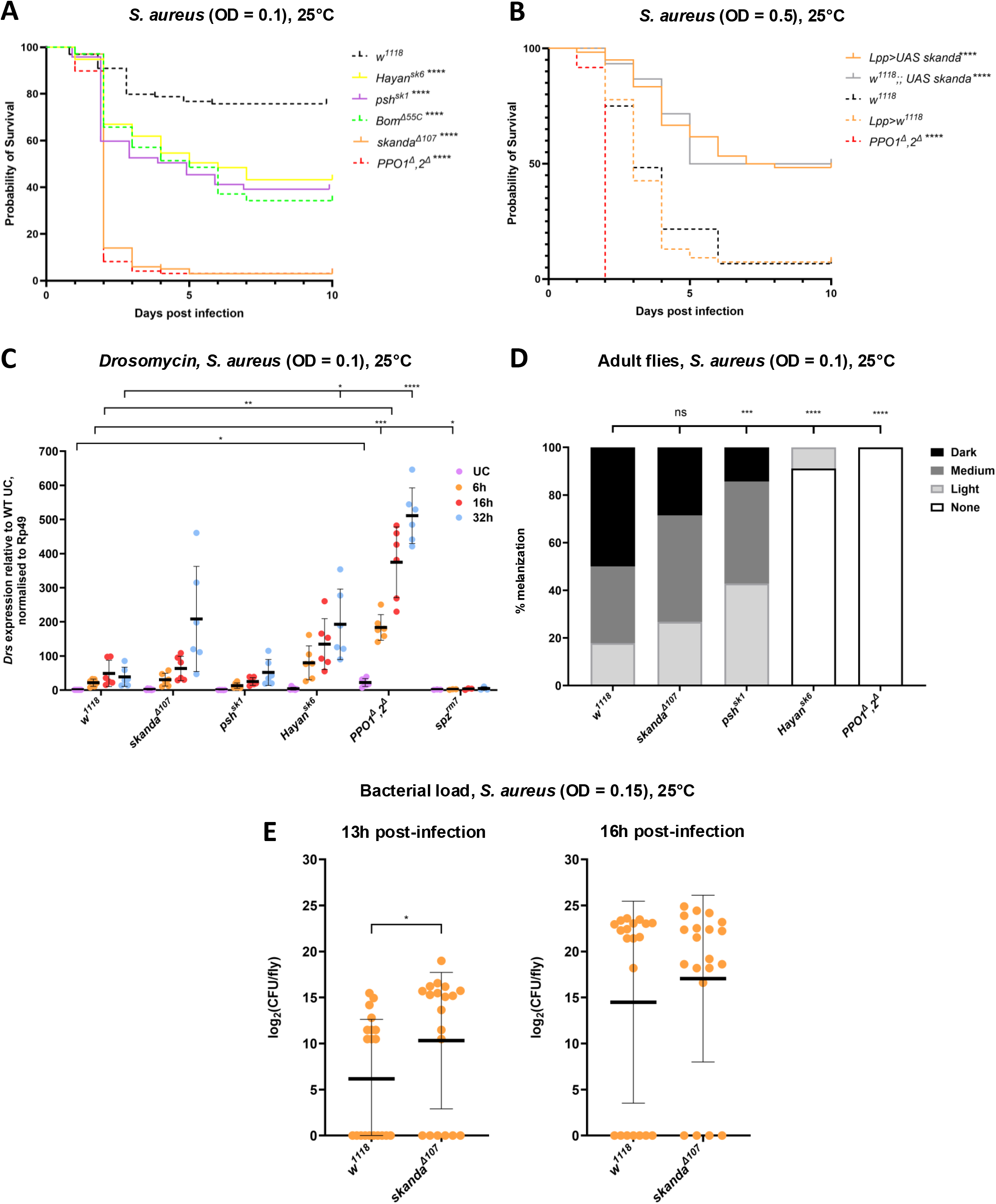
Skanda is critical for host defense to *S. aureus*. (A) Survival of *skanda* mutants was observed upon pricking with *S. aureus* (O.D.= 0.1). *Bom^Δ55C^* flies and *PPO1^Δ^,2^Δ^* flies were used as controls. *skanda^Δ107^*flies were highly susceptible to *S. aureus*, while *psh^sk1^*and *Hayan^sk6^* flies were moderately susceptible. (B) Survival of flies overexpressing *skanda* was observed upon pricking with *S. aureus* (O.D.= 0.5). *PPO1^Δ^,2^Δ^* flies were used as controls. Flies overexpressing *skanda* were more resistant to *S. aureus*. (C) *Drs* expression of *skanda* mutants was observed 6, 16 and 32 hours after pricking with *S. aureus* (O.D. =0.1). *PPO1^Δ^,2^Δ^* and *spz^rm7^* flies were used as controls. *skanda^Δ107^* and *Hayan^sk6^* flies showed increased *Drs* expression, while *psh^sk1^* flies showed wild-type *Drs* expression. *Drs* expression was strongly increased in *PPO1^Δ^,2^Δ^* flies and strongly reduced in *spz^rm7^* flies. Mixed-effects analysis indicated that there was a significant difference in *Drs* expression over time across genotypes (*p* < 0.0001 for both time and genotype factors. (D) Cuticular melanization of *skanda* mutants was observed 16 hours septic injury with a needle dipped in *S. aureus* OD_600_ 0.1. Dark, medium and light refers to the size and colour of the scab. *PPO1^Δ^,2^Δ^* flies were used as a positive control. *skanda^Δ107^* flies showed wild type melanization, while *psh^sk1^* flies showed reduced melanization. *Hayan^sk6^* showed strongly reduced melanization, while *PPO1^Δ^,2^Δ^* flies showed no melanization. (E) Bacterial load of *skanda* mutants was observed 24 hours post-pricking with *S. aureus* OD_600_ 0.1. *PPO1^Δ^,2^Δ^* and *spz^rm7^* flies were used as controls. *skanda^Δ107^*, *psh^sk1^* and *PPO1^Δ^,2^Δ^* flies showed increased bacterial load, while *Hayan^sk6^* and *spz^rm7^* flies showed wild-type bacterial load. ns = not significant (not shown for 6C, E), *p < 0.05, **p < 0.01, ***p < 0.001, and ****p < 0.0001.

To investigate the mode of host defense conferred by Skanda against *S. aureus*, we first checked *Drs* expression upon infection with a low dose of *S. aureus*. It should be noticed that the level of induction *Drs* upon systemic infection with a low dose of *S. aureus* was 50% lower than the level observed in flies infected with *M. luteus*, likely because of the low number of injected bacteria. Strikingly, *PPO1,2* mutants display a much higher expression of *Drs (***Figure 6C)**, a feature that has been associated to negative feedback of the melanization reaction on the Toll pathway (Liu et al., 2025). Although the biochemical analysis pointed to a negative role of Skanda on Psh and Hayan processing, *skanda^Δ107^* flies showed a wild-type Toll response as shown by the expression profile of Toll reporter genes *Drs* and *BomS1* upon *S. aureus* (**Figure 6C, Supp. Figure 6A**). Overexpressing *skanda* did not affect the Toll response to *S. aureus* (**Supp. Figure 6B**). The wild-type levels of *Drs* in *skanda^Δ107^* flies and *skanda* over-expressing flies rule out that the contribution of Skanda to host defense to *S. aureus* relies on the Toll pathway. We next checked the level of cuticular melanization upon septic injury with *S. aureus*. Septic injury with *S. aureus* induced an even stronger melanization response than that observed *M. luteus*, with an additional 10% of wild-type flies showing dark melanization. s*kanda^Δ107^* mutants once again had a wild-type cuticular melanization response, while *psh^sk1^* and *Hayan^sk6^*mutants had reduced cuticular melanization response (**Figure 6D**). A low residual cuticle blackening could be detected in *Hayan^sk6^*flies.

Overexpressing *skanda* using the *Lpp-Gal4* fat body driver led to a reduction in melanization to *S. aureus* as observed above for the experiment using clean injury (**Supp. Figure 6C**). Previous studies have shown that phagocytosis can contribute to the host defense against *S. aureus*. Several receptors, notably Eater and Draper, have been implicated in the phagocytosis of this bacterium (Bretscher et al., 2015; Melcarne et al., 2019; Shiratsuchi et al., 2012). We next investigated whether Skanda also contributes to phagocytosis of bacteria. We conducted an *ex vivo* phagocytosis assay using Alexa488-labeled *S. aureus* particles (Bretscher et al., 2015; Melcarne et al., 2019). Macrophages from third instar larvae were co-incubated with bacterial particles for 2 hours, and the phagocytic index was calculated. Bacterial phagocytosis was as wild-type in *skanda* mutant larval macrophages, indicating no overt role for Skanda in bacterial clearance (**Supp. Figure 6D**).

Survival to infection can be mediated by mechanisms that eliminate the pathogen (resistance mechanism) or by an increasing ability to endure the damage caused by the pathogen (disease tolerance) (Duneau et al., 2017; Soares et al., 2017; Westlake et al., 2024). Recent studies have shown a role for the Toll pathway in both resistance and tolerance [(Lou et al., 2026; Xu et al., 2022) but see (Tian et al., 2025)]. To distinguish between the two modalities, we monitored *S. aureus* load at 13 and 16 hours post infection with *S. aureus*, before *skanda* deficient flies were starting to die. Wild-type and s*kanda^Δ107^* flies showed a dichotomic bacterial count with flies having either high or low burden of *S. aureus* (**Figure 6E**). Such a pattern has already been observed, and reflects the stochasticity of the host defense with flies able or unable to control the bacteria (Duneau et al., 2017; Lafont et al., 2021; Ryckebusch et al., 2025). At the 13h timepoint, s*kanda^Δ107^*flies displayed a significantly higher bacterial burden compared to wild type, suggesting an early defect in pathogen control. This difference was no longer significant at 16h, indicating that the clearance gap between genotypes attenuates over time. The percentage of flies that have fully cleared the infection at 16 hours was however higher in the wild-type (8/20) than in s*kanda^Δ107^*(4/20). The fast killing induced by systemic *S. aureus* infection made difficult to find a condition that would allow to see a clear load difference Thus, while our result is consistent for a role of Skanda in resistance, Skanda could also contribute to tolerance to *S. aureus*.

We therefore conclude that the SPH Skanda contributes to host defense of *S. aureus* by a mechanism distinct from the Toll pathway, cuticular melanization and phagocytosis.

### Genetic dissection of the *skanda-psh-Hayan* gene cluster

The clustering of the three SP related genes *Hayan*, *psh* and *skanda* is suggestive of overlapping functions. The use of a short deletion *Hayan-Psh^Def^,* removing both genes, has recently revealed the partially redundant role of Hayan and Psh in integrating signals from both PRRs and microbial proteases that could not be revealed by single mutant analysis. Accordingly, flies lacking both *Hayan* and *psh* show a much lower level of Toll activation upon systemic infection with *M. luteus* when compared to flies lacking only one of the two genes (Dudzic et al., 2019). To better characterize the role of *skanda* in the context of the *Hayan-psh-skanda* gene cluster, we generated additional compound mutants. We first produced a short deletion *psh-skanda^Def^* that removed both *psh* and *Skanda*. We then used those flies to create, by *CRISPR-CAS9* targeted mutagenesis of *Hayan*, a *Hayan^2015^, psh-skanda^Def^* triple null mutant lacking the three genes (**Figure 5A**). Despite multiple attempts, we were unable to generate a *Hayan-skanda* double mutant. To circumvent this problem, we created flies carrying the *Hayan^2015^, psh-skanda^Def^* mutations with a genomic rescue of *psh* (referred to as *Hayan-skanda*^[*psh*+]^). We confirmed that *Hayan-skanda*^[*psh*+]^ flies had wild-type *psh* expression levels and that *psh-skanda^Def^* had no cis effects. We, however, observed reduced *skanda* expression in *Hayan-Psh^Def^* flies (**Supp. Figure 7A**). All the double and triple mutants were perfectly viable and displayed wild-type activation of the Imd pathway upon systemic infection with. Among all these double and triple mutants, only *Hayan^2015^, psh-skanda^Def^* and *Hayan-Psh^Def^* display a mild susceptibility to *Ecc15* (**Supp. Figure 7B and 7C**); consistent with a contribution to of the Toll pathway to combat this Gram-negative bacterium (Ryckebusch et al., 2025).

### *psh* and *skanda* redundantly regulate Toll signaling

We then explored the impact of double and triple mutations on three host mechanisms regulated by the Toll-PO SP cascade: i) Toll pathway activation, ii) cuticular melanization to wound and iii) survival to low dose of *S. aureus*.

Assessing *Drs* expression upon *M. luteus* expression and *B. subtilis* protease injection revealed a significant difference in *Drs* expression over time across genotypes. As previously reported, *Hayan-psh^Def^*flies have significantly reduced Toll pathway activity upon both systemic infection with *M. luteus* and injection of microbial proteases, confirming that both SPs redundantly regulate Toll signaling (**Figure 7A and 7B**). We however noticed a low level of residual *Drs* induction in *Hayan-psh^Def^* compared to *spz^rm7^*flies. *Hayan^2015^, psh-skanda^Def^* triple mutant flies also displayed a low level of *Drs* expression. Surprisingly, *psh-skanda^Def^*double mutant flies also displayed very low level of Toll pathway activity upon both *M. luteus* infection and injection of microbial proteases revealing a cryptic role of Skanda in Toll pathway activation in absence of *psh (***Compare 7A and 7B to Figure 5B and 5C**). Finally, *Hayan-skanda*^[*psh*+]^ double mutant flies exhibited a wild-type level of activation of Toll pathway activation upon *M. luteus* and upon injection of *B. subtilis* proteases (**Figure 7A and 7B**). Collectively, the use of double mutants reveals a role for Skanda in the regulation of the Toll pathway along with Hayan and Psh. While a mutation in Hayan in the absence or presence of Skanda has little impact on Toll pathway activity, the loss of Psh has a much stronger impact on the Toll pathway activity in absence of either Skanda or Hayan.

**Figure 7.**
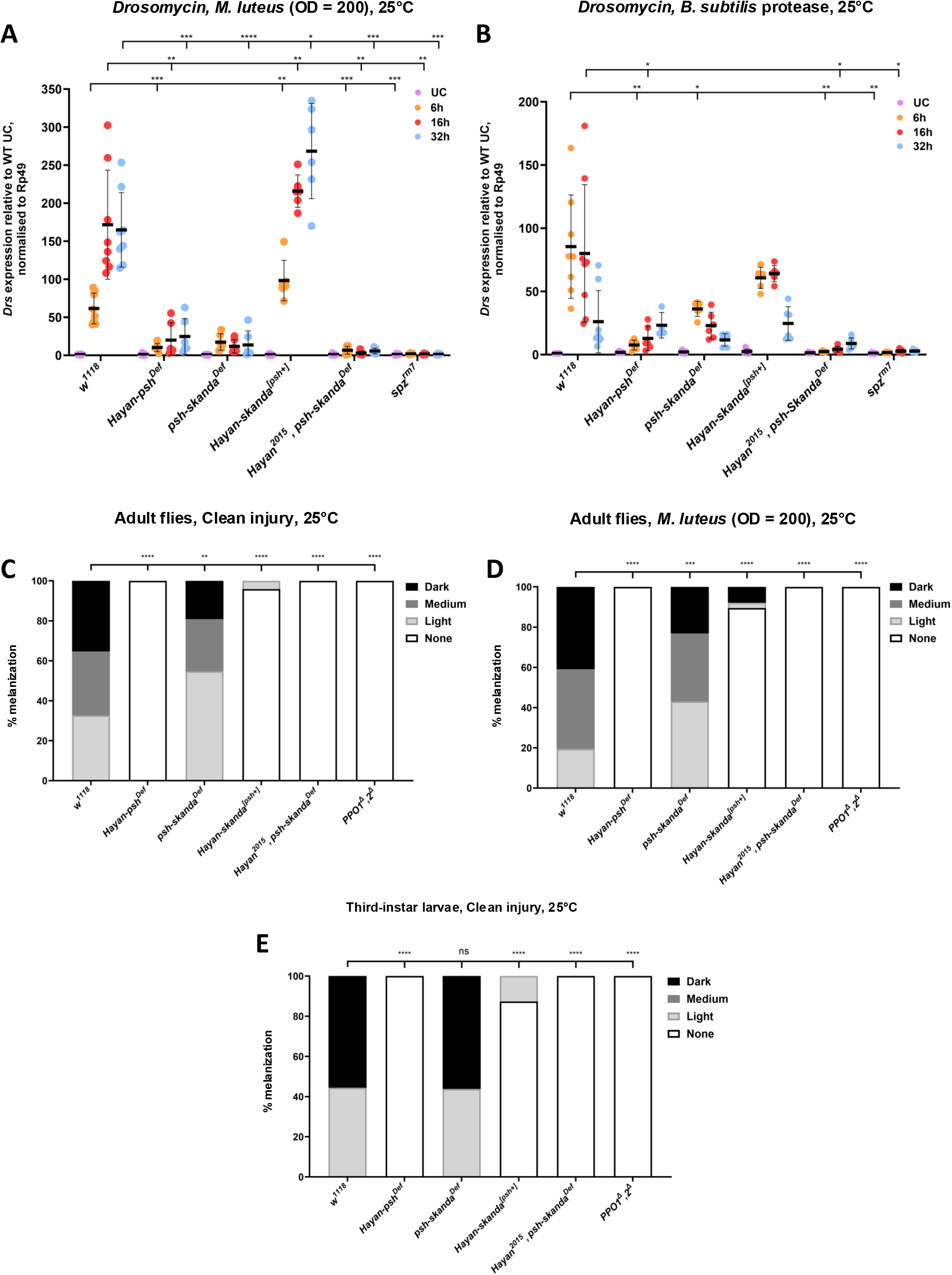
Skanda and Psh redundantly regulate the Toll pathway. (**A,B**) *Drs* expression of compound mutants of the *skanda-psh-Hayan* cluster was observed 6, 24 and 48 hours after **(A)** pricking with *M. luteus* OD_600_ 200 and **(B)** injection with *B. subtilis* protease. *spz^rm7^* flies were used as a positive control. *Hayan-psh^Def^*, *psh-skanda^Def^*, *Hayan^2015^, psh-skanda^Def^* and *spz^rm7^* flies showed strongly reduced *Drs* expression, while *Hayan-skanda^[psh+]^* flies showed wild-type *Drs* expression. (**C,D**) Cuticular melanization of *Hayan-psh-skanda* compound mutants was observed 16 hours post clean injury with a sterile needle **(C)** and septic injury with a needle dipped in *M. luteus* (O.D.= 200 **(D)** Dark, medium and light refers to the size and colour of the scab. *PPO1^Δ^,2^Δ^* flies were used as a positive control. *psh-skanda^Def^* flies showed slightly reduced melanization, while *Hayan-skanda^[psh+]^* flies showed strongly reduced melanization. *Hayan-psh^Def^*, *Hayan^2015^, psh-skanda^Def^* and *PPO1^Δ^,2^Δ^*flies showed no melanization. (**E**) Melanization of compound mutant third instar larvae of the *skanda-psh-Hayan* cluster was observed 2 hours post clean injury with a sterile needle. *PPO1^Δ^,2^Δ^* larvae were used as a positive control. *psh-skanda^Def^* larvae showed wild-type melanization, while *Hayan-skanda^[psh+]^* flies showed strongly reduced melanization. *Hayan-psh^Def^*, *Hayan^2015^, psh-skanda^Def^* and *PPO1^Δ^,2^Δ^*larvae showed no melanization. Although presented separately for clarity, all experiments with double and triple mutants shown in this figure and 8 were carried out concurrently with those in Figures 5 and 6, enabling direct comparison between single and compound mutants ns = not significant (not shown for 7A, B), *p < 0.05, **p < 0.01, ***p < 0.001, and ****p < 0.0001.

### Loss of *skanda* does not further enhance the cuticular melanization defects caused by the loss of *Hayan* or *psh*

We next monitored the impact of double and triple mutations on cuticular melanization upon clean injury and septic injury with *M. luteus* of adult flies as described above. *psh-skanda^Def^* flies had a slightly weaker overall level of melanization to clean injury when compared to wild-type flies (**Figure 7C**), like *psh* single mutants (**Figure 5D**). *Hayan-psh^Def^*, *Hayan-skanda*^[*psh*+]^ double mutant and *Hayan^2015^, psh-skanda^Def^* triple mutant flies showed no melanization upon clean injury, similar to *Hayan* single mutant flies (**Compare Figure 7C to 5C**). Analyzing melanization at the wound site upon septic injury with *M. luteus* or upon pricking of mutant L3 larvae yielded similar results (**Figure 7D and 7E**). This experiment confirms that Hayan is the main executive serine protease that regulates cuticular melanization, with a minor contribution from Psh. The absence of Skanda did not further impact melanization in absence of *psh* or *Hayan*.

### *Hayan-psh-skanda* compound mutants are also susceptible to *S. aureus*

The use of single mutant flies revealed a critical role of Skanda in the survival to low dose of *S. aureus* with *skanda* deficient flies reaching the same degree of susceptibility as *PPO1,2* double mutants. In contrast, loss of *psh* or *Hayan* single mutant lead to an intermediate susceptibility phenotype similar to that observed with *Bom55C* mutants (and *spz^rm7^* see (Dudzic et al., 2019)). We then explored whether the absence of two or three SPs of the *skanda-psh-Hayan-*gene cluster affects host defense to this bacterium. We observed that all compound mutants were highly susceptible to infection with low dose of *S. aureus* (**Figure 8A**). The absence of *Hayan* or *psh* did not aggravate the susceptibility of *skanda* flies to *S. aureus*, albeit a slightly better survival was observed in *psh*, *skanda* double mutant compared to *skanda* single mutant (**compare Figure 8A to 6A**). As previously reported, we observed that the *psh*,*Hayan* double mutants display a marked susceptibility to *S. aureus*, compared to *psh* and *Hayan* single mutants, revealing that both SPs redundantly regulate activity against *S. aureus*.

**Figure 8.**
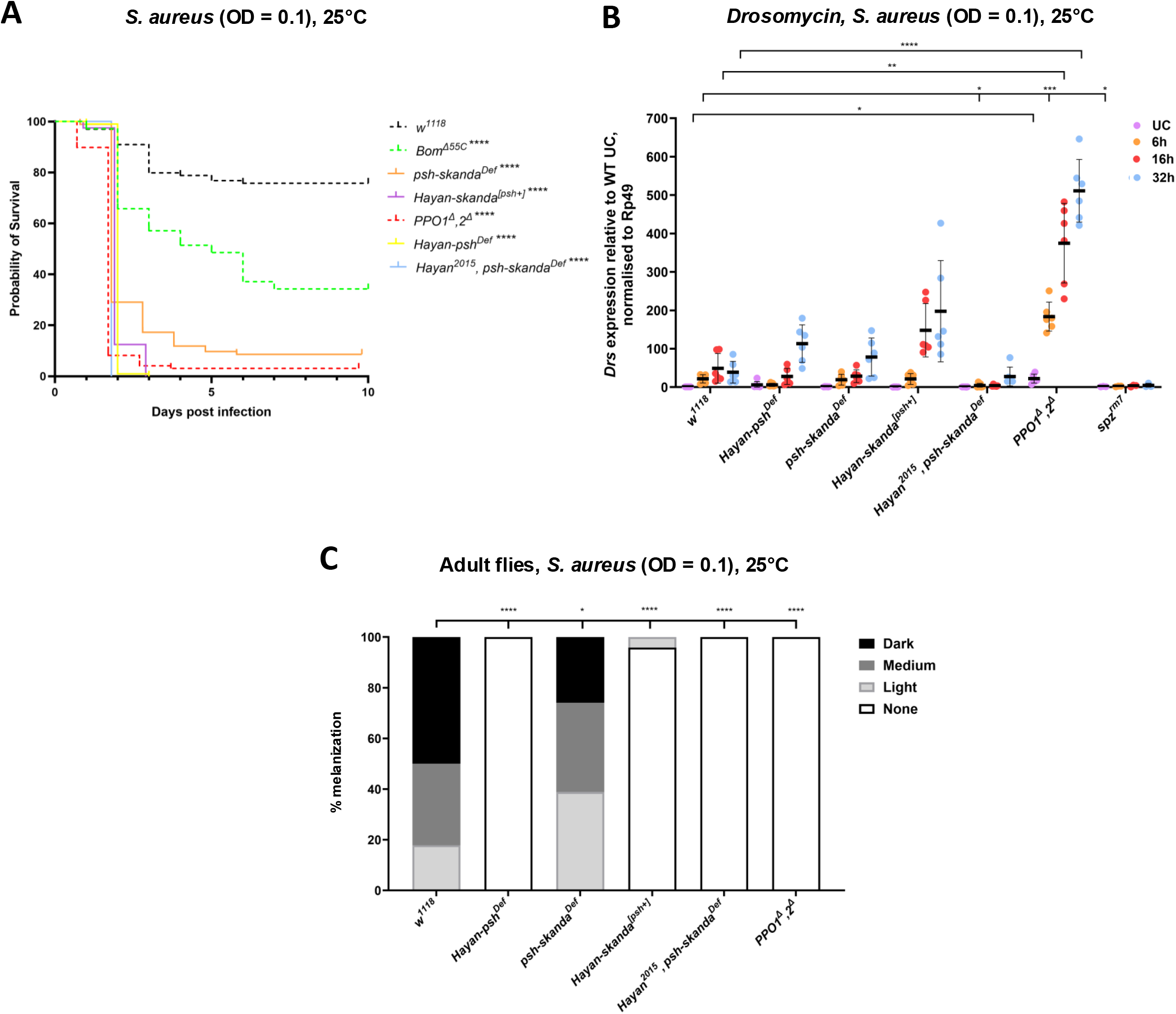
*Hayan-psh-skanda* compound mutants are susceptible to *S. aureus* infection. (**A**) Survival of *Hayan-psh-skanda* compound mutants was observed upon pricking with *S. aureus* (O.D.= 0.1). *Bom^Δ55C^* flies and *PPO1^Δ^,2^Δ^*flies were used as controls. All compound mutants were highly susceptible to *S. aureus*. (**B**) *Drs* expression of *Hayan-psh-skanda* compound mutants was observed 6, 16 and 32 hours after pricking with *S. aureus* OD_600_ 0.1. *PPO1^Δ^,2^Δ^* and *spz^rm7^* flies were used as controls. All compound mutants showed increased *Drs* expression 32 hours post infection. *Drs* expression was strongly increased in *PPO1^Δ^,2^Δ^* flies and strongly reduced in *spz^rm7^* flies. Mixed-effects analysis indicated that there was a significant difference in *Drs* expression over time across genotypes (*p* < 0.0001 for both time and genotype factors). (**C**) Cuticular melanization of *Hayan-psh-skanda* compound mutants was observed 16 hours septic injury with a needle dipped in *S. aureus* OD_600_ 0.1. Dark, medium and light refers to the size and colour of the scab. *PPO1^Δ^,2^Δ^*flies were used as a positive control. *psh-skanda^Def^* flies showed reduced melanization, while *Hayan-skanda^[psh+]^* flies showed strongly reduced melanization. *Hayan-psh^Def^*, *Hayan^2015^, psh-skanda^Def^* and *PPO1^Δ^,2^Δ^* flies showed no melanization. ns = not significant (not shown for 8B). *p < 0.05, **p < 0.01, ***p < 0.001, and ****p < 0.0001.

We next explored the level of Toll pathway activation and cuticular melanization upon systemic infection with a low dose of *S. aureus*. The *Drs* expression profile of these mutants also revealed something unexpected. Experiments reported by Dudzic at al., 2019 and here above showed that *Hayan-psh^Def^*and *psh-skanda^Def^* double mutant flies show almost no *Drs* expression upon systemic infection with *M. luteus*.

However, when pricked with *S. aureu*s, they show a much higher expression of *Drs*, similar to the expression levels of *psh^sk1^*flies (**Figure 8B**). Additionally, all compound mutants showed a peak in expression 32 hours post-infection, which was not observed in *spz^rm7^* mutants. This residual level of Drs expression indicates that an SP distinct from Hayan and Psh must be capable of activating SPE or another Spätzle processing enzyme upon *S. aureus* infection as suggested by (Yamamoto-Hino and Goto, 2016). Analysis of cuticular melanization upon systemic infection with *S. aureus* confirms the central role of Hayan in controlling cuticular melanization downstream of the Toll-PO cascade (**Figure 8C**). Collectively, the use of compound mutants confirms the major role of *skanda* in host defense to *S. aureus* but also points to the existence of a serine protease in addition to Hayan and Psh regulating the processing of the Toll ligand Spätzle.

## Discussion

Serine proteases and their noncatalytic homologs are critical regulators of insect immune responses, particularly in the activation of the Toll signaling pathway and melanization (Kanost and Jiang, 2015; Veillard et al., 2016). While genetic studies in *Drosophila* have identified many components of the SP-SPH system, recent biochemical work has begun to elucidate the underlying mechanisms in greater detail. Notably, *in vitro* reconstitution of the SP cascade has provided a robust biochemical framework that supports over three decades of genetic findings (Shan et al., 2023). This work also led to the discovery of a previously uncharacterized gene, *cSP48*, and revealed an unexpected function for the protein Necrotic, which blocks the activity of Grass—thus opening new directions for genetic validation. In a separate investigation, combining biochemical assays and reverse genetics, Jin et al. (2023) showed that cSPH35 and cSPH242 form a cofactor complex that significantly enhances proPO1 activation by MP2 in *Drosophila* (Jin et al., 2023). This cofactor mechanism appears to be evolutionarily conserved among holometabolous insects, including mosquitoes, lepidopterans, and beetles (Jin et al., 2023; Park et al., 2011; Wang et al., 2025, 2020). However, as more biochemical data emerge to complement genetic analyses, discrepancies between *in vitro* and *in vivo* findings are inevitable, due to the distinct strengths and limitations of each approach. Although *in vitro* assays are well-controlled, and highly reproducible, their simplification can lead to a lack of physiological complexity observed in whole-organism studies, limiting their ability to uncover certain regulatory mechanisms. In contrast, genetic *in vivo* approach, although powerful to identify critical components of a pathway can miss important but redundant interactions and is limited in establishing direct interactions. In this context, we report the biochemical and genetical characterization of the serine protease homologue Skanda. Although discrepancies are observed between the two approaches, both the biochemical and genetic studies reveal an important role of this SPH in Toll-PO SP mediated immunity.

Skanda exhibits several distinctive biochemical properties: (i) the presence of two clip domains, (ii) the absence of a linkage between the interdomain Cys pair, distinguishing Skanda from cSPH35 and cSPH242; (iii) the presence of two unique disulfide bonds that stabilize the PLD domain further; and (iv) a 81-residue region rich in *O*-glycosylated Ser and Thr. These characteristics may hold the key to understanding the specific role of Skanda. Our study shows that Skanda is an unstable protein and can be further processed by Grass and cSP48 of the Toll-PO SP cascade. Our biochemical analysis did not reveal any impact of Skanda on the processing of upstream components of the Toll-PO SP cascades such as Grass or cSP48. Thus, under *in vitro* conditions, Skanda behaves like a placeholder to reduce the proteolytic activation of Psh and Hayan and their downstream effectors that must be processed Spz and PPO. This placeholder is consistent with the genomic clustering and high similarity of *skanda* with *psh* and *Hayan*. We speculate that a protein factor or a surface feature, through an interaction with the ST-rich region in Skanda, may increase the cleavage of Skanda in the clip domains *in vivo* and, through clip domain interaction, modulate the production of Psh, Hayan, Sp7, SPE, MP1, Ser7, and consequently mature Spätzle, PPO1, and PPO2. Insertion of a similar ST-rich region in *Anopheles gambiae* CLIPA7f greatly reduced the cofactor activity of CLIPA7s when paired with CLIPA4 (Wang et al., 2025). This regulation, which remains to be identified, might act a molecular switch that modulates Skanda’s function, elevating our understanding of system-level regulation. Thus, the biochemical approach points to a role of Skanda in reducing Toll-PO SP activation. While, we favor a role of Skanda at the Psh-Hayan level, we cannot exclude that this SPH modulates an SP working upstream in the Toll-PO cascade, establishing a negative feedback loop. It is well established that serine proteases must be tightly regulated to avoid excessive, deleterious activation. Mutations in serpin genes encoding Spn27A Spn28D, Spn77B, Spn5, or Necrotic that negatively regulate the Toll-PO SP cascades and lead to constitutive activation of the Toll and/or melanization pathway cause severe lethality (Ahmad et al., 2009; E. De Gregorio et al., 2002a; Levashina, 1999; Scherfer et al., 2008, p. 77; Tang et al., 2008). In addition to serpins, serine protease homologs such as Skanda might modulate these cascades in a more subtle way, adjusting the level of immune reactivity.

Although the biochemical study suggests a negative role for Skanda, the genetic approach reveals a more complex picture. Use of loss-of-function mutation shows that Skanda does not repress Toll pathway signaling upon systemic infection with *M. luteus*, *S. aureus* or injection of microbial protease. However, Skanda redundantly positively regulates the Toll pathway with Psh since *psh,skanda* double mutants show a much lower Toll signaling activity upon *M. luteus* infection than *psh* single mutant flies. We have currently no hypothesis to explain the difference between the biochemical and genetic approaches, although we cannot exclude that Sf9 cells used in the biochemical analysis produce glycosylation patterns of Skanda that are different from *Drosophila* ones. Alternatively, the inhibitory or stimulatory role of Skanda on Psh and Hayan could depend of the stoichiometry of these proteins that likely differs in the in vitro and in vivo approaches.

We also observed that Skanda has little impact on cuticular melanization which is mainly mediated by Hayan, but its over-expression reduced melanization. This latter result could reflect an artefact of overexpression. Nevertheless, it supports the biochemical analyses, pointing to a negative role of Skanda in cuticular melanization. However, the most striking phenotype of *Skanda* mutation is the high susceptibility to *S. aureus* infection. Thus, Skanda non-redundantly contributes to producing microbicidal activity against *S. aureus*, which is distinct from cuticular melanization and Toll pathway activation. To date, we have little information on the nature of this microbicidal activity, although a role of ROS produced during the melanization reaction has been suggested (Westlake et al., 2024). However, we know that it also involves Sp7, PPO1 and the Hayan-Psh platform. In contrast to mutations affecting these factors that all reduce melanization at the wound site, Skanda display a much more specific phenotype. At this stage, we cannot propose a mechanism of action of Skanda as we lack knowledge of the factors that link the Toll-PO SP cascade to this microbicidal activity. Since Skanda most likely has a regulatory function, we can speculate that Skanda acts at the level of Psh-Hayan to allow Hayan to activate the Toll pathway. Skanda would skew the activity of the Psh-Hayan platform to induce Toll signaling and host defense to *S. aureus* rather than cuticular melanization. Such a hypothesis would explain why *psh, Skanda* double mutants fail to activate the Toll pathway and why over-expression of Skanda tend to reduce cuticular melanization. In addition, Skanda would be critical for the Hayan-Psh platform to activate the killing of *S. aureus*, possibly by favoring the cleavage of Sp7 and/or PPO1. Thus, Skanda would tailor the Toll-PO SP cascade to more specific immune functions; limiting cuticular melanization and favoring microbicidal activity. This model is consistent with the biochemical observation that Skanda can reduce the cleavage of Hayan but is in disagreement with the observation that this SPH can slightly lessen Psh cleavage. The presence of Skanda in the hemolymph (Rommelaere et al., 2025) suggests a role in the systemic immune response; however, we cannot exclude that it may be particularly important within the local microenvironments at sites of injury. Future studies should be done to reconcile the biochemical and genetical approaches and better characterize the important steps downstream of Psh and Hayan.

Overall, our study identified a new regulatory serine protease homolog of the Toll-PO SP cascade, but also underlines how far we are from understanding the whole complexity of these cascades that shape the systemic immune response of insects.

## Materials and Methods

### Key Resources

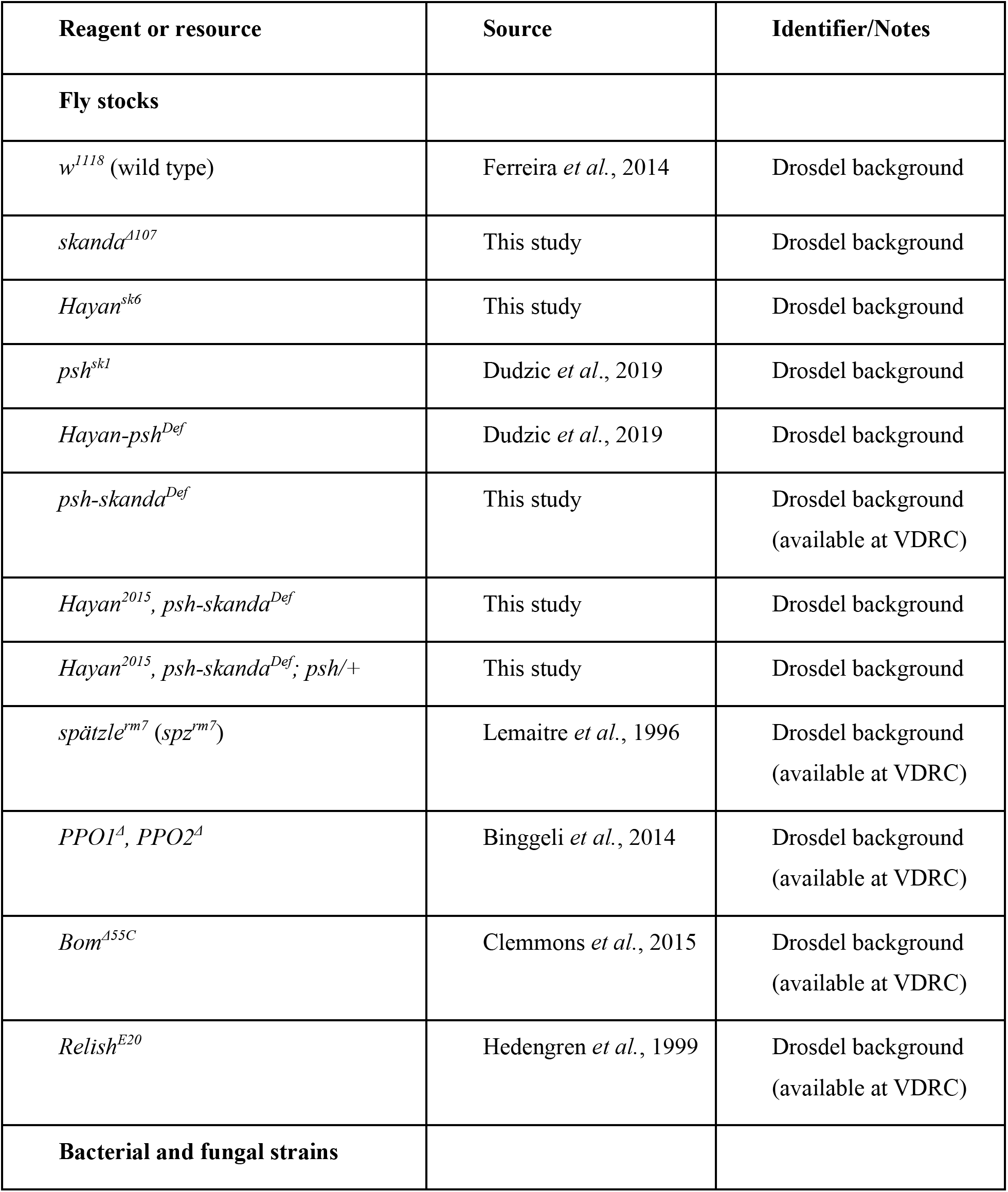
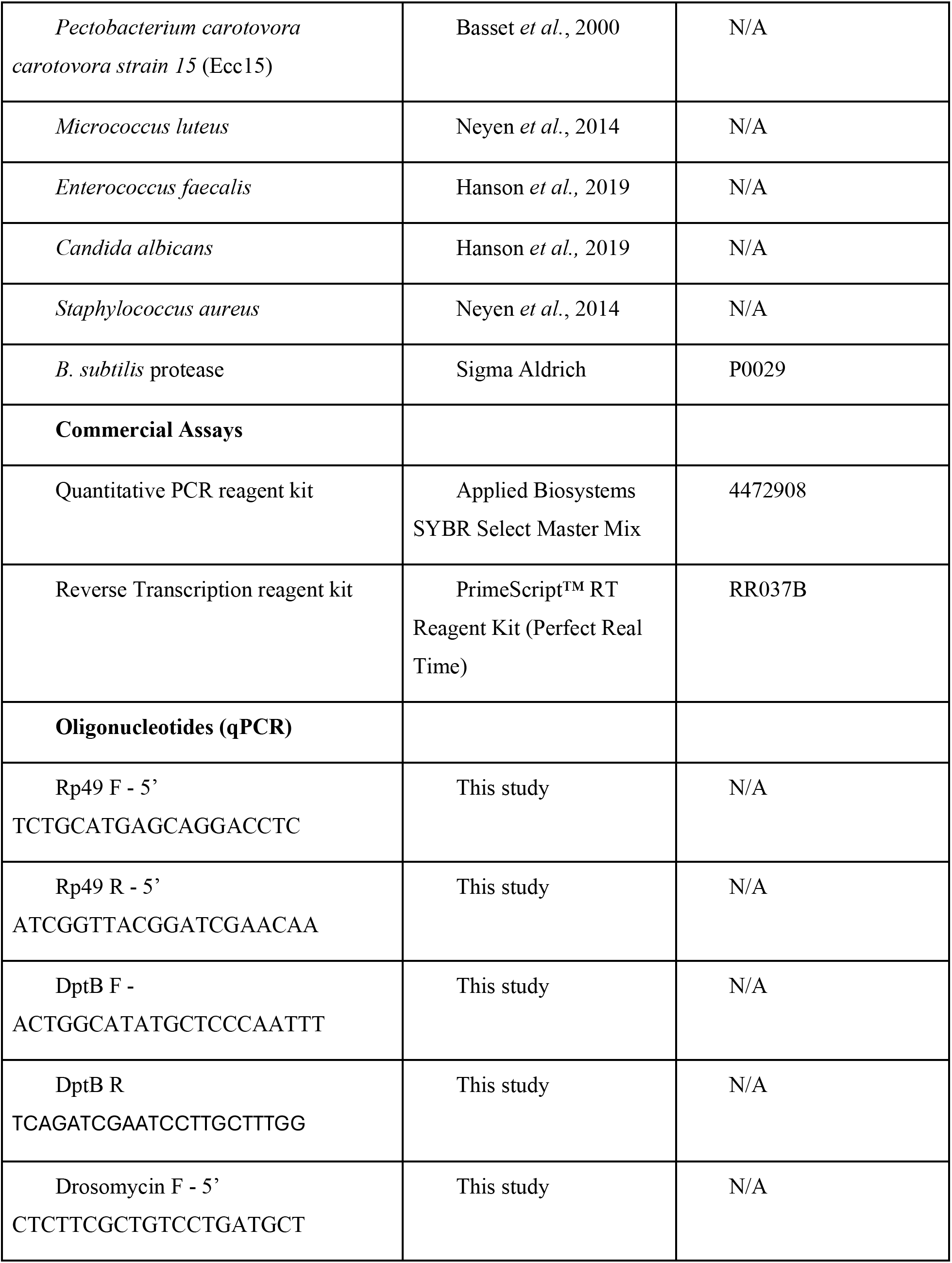
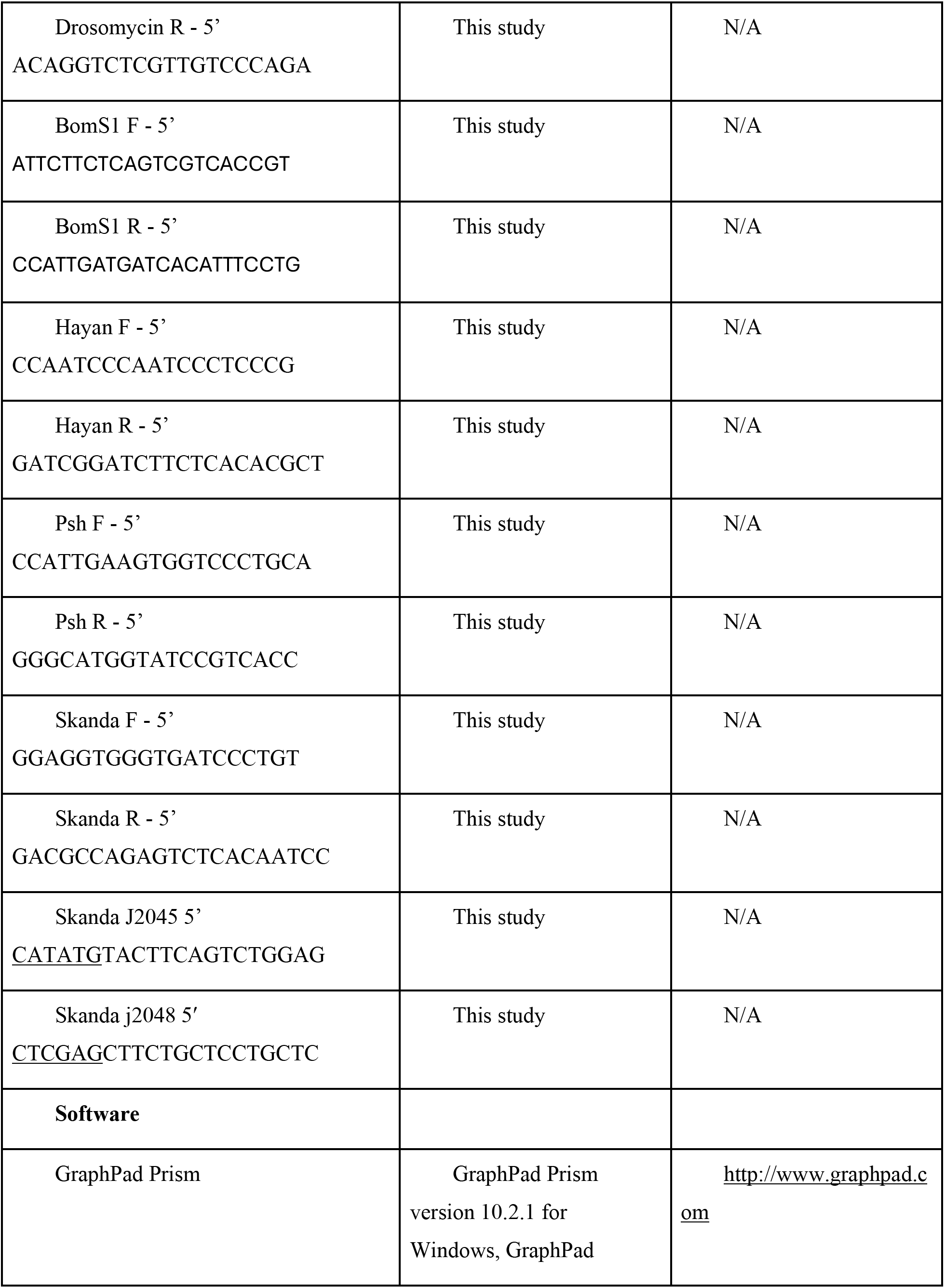
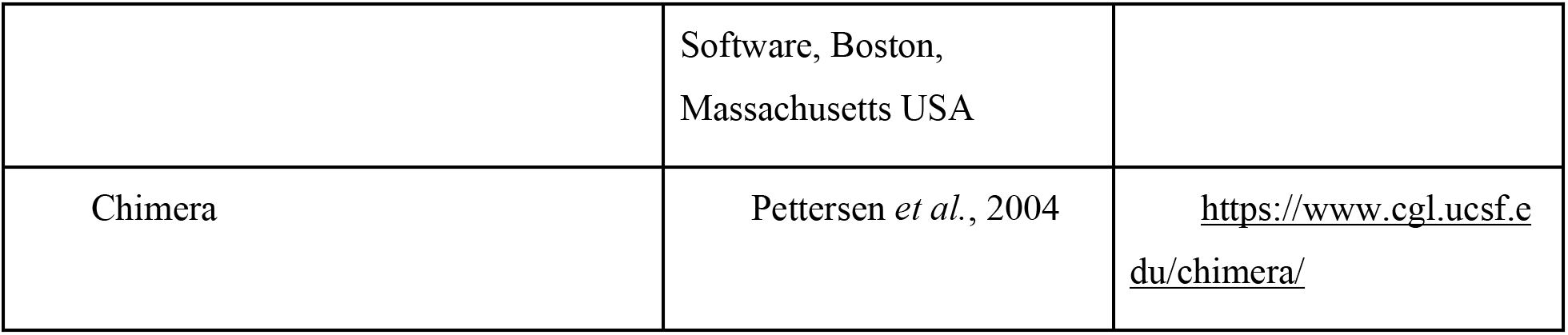

### Cloning and production of *D. melanogaster* Skanda

The coding region of mature proSkanda was amplified by PCR using primer pairs j2045 (5′-<u>CATATG</u>TACTTCAGTCTGGAG) and j2048 (5′-<u>CTCGAG</u>CTTCTGCTCCTGCTC), with the cDNA clone X (DGRC) as the template. The resulting 1,419 bp PCR product was cloned into the pGEM-T vector (Promega) and sequence verified. The *Nde*I-*Xho*I fragment encoding full-length Skanda was excised and ligated into the pMHFFH6 expression vector (Shan et al., 2023). The recombinant Skanda includes a coding region flanked by an N-terminal hemagglutinin (HA) tag and a C-terminal FLAG–6×His tag, with the final sequence: GIHYPYDVPDYAHMY^26^FS…EQK^494^LEDYKDDDDKHHHHHH. A recombinant baculovirus stock (1–2×10⁸ pfu/ml) was generated using this construct and used to infect *Spodoptera frugiperda* (Sf9) cells at a density of 2.4×10⁶ cells/ml in 1,000 ml of Sf-900™ III serum-free medium, at a multiplicity of infection (MOI) of 5–10. Following incubation at 27 °C for 96 h with gentle agitation, conditioned medium was clarified by centrifugation, and Skanda was purified from the supernatant by ion exchange chromatography on a dextran sulfate-Sepharose column (Jin et al., 2024). Fractions containing Skanda were identified via immunoblotting, pooled, and subjected to further purification on a 1.0 ml Ni²⁺-nitrilotriacetic acid (Ni-NTA) agarose column. Bound proteins were eluted with a 0–0.25 M imidazole gradient. The eluted Skanda was concentrated and buffer exchanged to 20 mM Tris-HCl, 50 mM NaCl (pH 7.5) using a 30 kDa molecular weight cut-off centrifugal filter device (Sigma-Aldrich). Purified aliquots were flash-frozen in liquid nitrogen and stored at -80 °C.

### Preparation of *D. melanogaster* serine protease zymogens

Precursors of *Drosophila* serine proteases including ModSP, SP48, Grass, Persephone, Hayan-PA, Hayan-PB, Sp7, SPE, MP1, Ser7, and Spätzle, as well as prophenoloxidase-1 (proPO1) were either obtained from previous preparations (Shan et al., 2023) or freshly expressed and purified following identical protocols when stocks were depleted. Notably, a substantial portion of proModSP underwent spontaneous activation during expression, enabling it to initiate the downstream protease cascade in the absence of microbial surface polysaccharides or their recognition proteins. Information about the recombinant proteins and their antibodies is provided in Table S2.

### Identification of Skanda cleavage sites by LC-MS/MS analysis

To identify proteolytic cleavage sites in Skanda, purified Skanda (1 μg) was incubated with ModSP (0.5 μg), proSP48 (0.1 μg), and proGrass (0.4 μg) in 25 μl of reaction buffer A (20 mM Tris-HCl, pH 7.5, 5 mM CaCl_2_, 0.001% Tween-20) at 37 °C for 1 h. Control reactions included Skanda alone and combinations omitting individual proteases. After SDS-PAGE, gel bands corresponding to ∼85, 44, and 35 kDa were excised based on pre-stained protein markers. Samples were reduced with tris(2-carboxyethyl)phosphine (TCEP), alkylated with iodoacetamide, and digested with trypsin or chymotrypsin. LC-MS/MS was performed as previously described (He et al., 2016) Raw data were searched using Byonic (v3.5.3) with default parameters unless otherwise specified. Arg or Lys was specified as a cleavage site, and up to two missed cleavages are allowed. The precursor mass tolerance was 15 ppm, and the fragment mass tolerance was 0.5 Da. All modifications were treated as variables, including carbamidomethylation and propionamide on Cys residues, oxidation of His, Met and Trp, N-terminal acetylation or pyro-Glu formation. Spectra were searched against a *Drosophila* database, with decoys and contaminant sequences appended. A protein-level false discovery rate of 1% was applied. Peptides from Skanda, identified in the Byonic search with PEP2D < 0.001 were selected. Extracted ion chromatograms were generated in Qual Browser with mass tolerance of 5 ppm and Genesis for peak detection. The peak area corresponding to the consistent scan time was taken as the peptide intensity. A total of 19 peptides from Skanda were retained for quantification.

### Insect stocks

Fly stocks were raised at 25°C on Yeast-Cornmeal food (6% cornmeal, 6% yeast, 0.62% agar, 0.1% fruit juice, supplemented with 10.6g/L Moldex and 4.9ml/L propionic acid). Experiments were performed on 5 to 10-day-old male flies. *w^1118^* Drosdel flies were used as wild type unless stated otherwise. The *psh^sk1^*, *Hayan-psh^Def^*, *spätzle^rm7^*, *PPO1^Δ^, PPO2^Δ^*, *Bom^Δ55C^* and *Relish^E20^* lines are described previously (Lemaitre *et al*., 1996; Hedengren *et al*., 1999; Binggeli *et al*., 2014; Clemmons *et al*., 2015; Dudzic *et al*., 2019). The *Hayan^sk6^* and *psh-skanda^Def^*mutant lines were generated using CRISPR/Cas9 as described in (Kondo and Ueda, 2013). The *Hayan^sk6^* line has a 14 bp deletion from position 18482430 to 18482443 on the *X* chromosome. The *psh-skanda^Def^* line has a 4623 bp deletion from position 18484829 to 18489451 on the *X* chromosome. The *skanda^Δ107^* and *Hayan^2015^, Psh-Skanda^Def^* mutant lines were generated using CRISPR/Cas9 as described in (Rommelaere et al., 2019). The *skanda^Δ107^* line contains an 8 bp deletion from position 18488965 to 18488972 on the X chromosome. The *Hayan^2015^, psh-skanda^Def^* line was created using the *psh-skanda^Def^* line, and contains an additional 5 bp deletion from position 18482349 to 18482353 on the X chromosome. The Skanda cDNA was first cloned into the pENTR entry vector and then transferred into the UASg-attB vector. The final UASg-attB-Skanda construct was then inserted into flies at the attP2 landing site on the third chromosome. All the mutant strains were isogenised in the *w^1118^* Drosdel background. To create the *Hayan-skanda* double mutant, the *psh* genomic locus was inserted into an Attp40 site on the second chromosome. This line was then crossed with *Hayan^2015^, psh-skanda^Def^* flies to create a genomic rescue of *psh*.

### Microorganism infection experiments

The bacterial strains and the optical densities of their pellets at 600 nm (OD_600_) were: *Pectobacterium carotovora carotovora strain 15* (Ecc15, OD_600_ 200), *Micrococcus luteus* (OD_600_ 200), *Candida albicans* (OD_600_ 200); *E. feacalis* OD_600_ 5 and *Staphylococcus aureus* (*S. aureus*, OD_600_ 0.1, OD_600_ 0.15 and OD_600_ 0.5). All strains were cultured in Luria Broth (LB) overnight. *M. luteus* and *S. aureus* were cultured at 37°C, while Ecc15 was cultured at 29°C. Pellets were diluted in 1X PBS. Systemic infection (septic injury) experiments were performed by pricking adults in the thorax with a thin needle of diameter 0.2mm previously dipped into a concentrated bacterial pellet.

### Survival analysis and bacterial load quantification

5-10 days old flies were used for survival experiments that were performed either at 25°C (*S. aureus*, OD_600_ 0.1 or 0.5) or 29°C (*Ecc15*, OD_600_ 200, *C. albicans*). 20 flies per genotype were used in each experiment and survival was scored once daily. Three independent survival experiments were conducted per microorganism.

5-10 days old flies were used for bacterial load quantification. Flies infected with *S. aureus* (OD_600_ 0.15) were maintained at 25°C and collected at indicated time points post-infection. They were surface sterilized by washing them in 70% ethanol. Ethanol was removed, and flies were homogenized using a Precellys bead beater at 6500 rpm for 30 s in LB broth with 100 μL added per fly. These lysates were serially diluted and 100 µL was plated on LB agar. Plates were incubated overnight, and colony-forming units (CFUs) were counted manually.

### Quantitative RT-PCR

Infected flies at 25°C (*M. luteus,* OD_600_ 200, *B. subtilis* protease 1:2000, and *S. aureus*, OD_600_ 0.1) or 29°C (*Ecc15*, OD_600_ 200) were collected at indicated time points post-infection along with unchallenged controls and homogenized using a Precellys bead beater at 10000 rpm for 2 cycles of 30 s each. Total RNA was isolated using TRIzol reagent and dissolved in DEPC-treated RNase-free water. 500 ng of total RNA was reverse-transcribed in 10 µL reactions using PrimeScript RT (TAKARA) with random hexamer and oligo dT primers to selectively amplify mRNA. Quantitative PCR was performed on a LightCycler 480 (Roche) in 96-well plates using the Applied Biosystems SYBR Select Master Mix. Three independent experiments were conducted per microorganism, with 2 replicates of 5 flies per genotype each time.

### Melanization assay

The thorax of adult flies was pricked using a 0.2 mm needle previously dipped into ethanol (clean injury) or a concentrated bacterial pellet (*M. luteus*, OD_600_ 200 and *S. aureus*, OD_600_ 0.1) and maintained at 25°C. Pictures of melanized flies were taken 16 hours post-injury, using a Leica M 205 FA microscope, a Leica DFC7000FT camera and the Leica Application Suite. Four independent experiments were conducted using 20 flies per genotype each time. Third-instar larvae were pricked in the hind-region using an ethanol-sterilized needle of diameter 0.1 mm and maintained at room temperature. Intensity of melanization was assessed 2 hours post-injury according to criteria described in Supplementary Figure 5.

### *Ex vivo* larval hemocyte phagocytosis assay

Five third instar larvae were surface-sterilized and bled into 120 μL of Schneider’s medium within 2 minutes. The hemolymph was collected in Protein LoBind Eppendorf tubes. 105 bacterial bioparticles were added to the hemocyte suspension and incubated at room temperature for 1 hour. After incubation, the samples were placed on ice, and hemocytes were immediately analyzed using a CytoFLEX flow cytometer (Beckman Coulter).

The phagocytic index (PI) was calculated using the following equations:

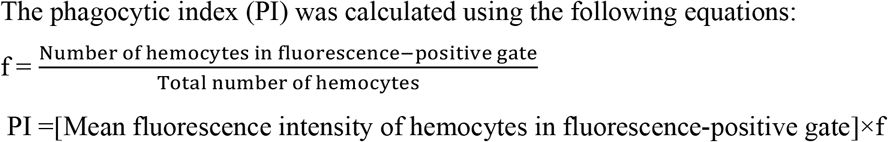

### Statistical analysis

The statistical significance of survival data was calculated using a log-rank test compared to wild-type flies. RT-qPCR data from bacterial infection and protease injection was analyzed using Mixed-effect analysis with Dunnett’s multiple comparisons. The blackening strength (dark, medium, or light) of the adult cuticle was analyzed using Chi-square test. Bacterial load was compared using the Mann-Whitney test. RT-qPCR data to assess cis-effects of mutations and UAS-expression, and phagocytosis data was analyzed using one-way ANOVA with Dunnett’s multiple comparisons. *p*values of < 0.05 = *, < 0.01 = **, < 0.001 = ***, and < 0.0001 = **** were considered significant.

Graphs were plotted using GraphPad Prism.

## Acknowledgments

We thank Samuel Rommelaere and Prince Kumar Sah for experimental help. This project was supported by the SNSF grants (310030_215073).

## Author Contributions

Conceived and designed the experiments: SV, AK, HJ, BL. Performed the experiments: SV,YW,CX, TS. Analyzed the data: Contributed reagents/materials/analysis tools: J-PB. Wrote the paper: SV, HJ, BL.

## Supplementary figures

**Supplementary Figure 1.**
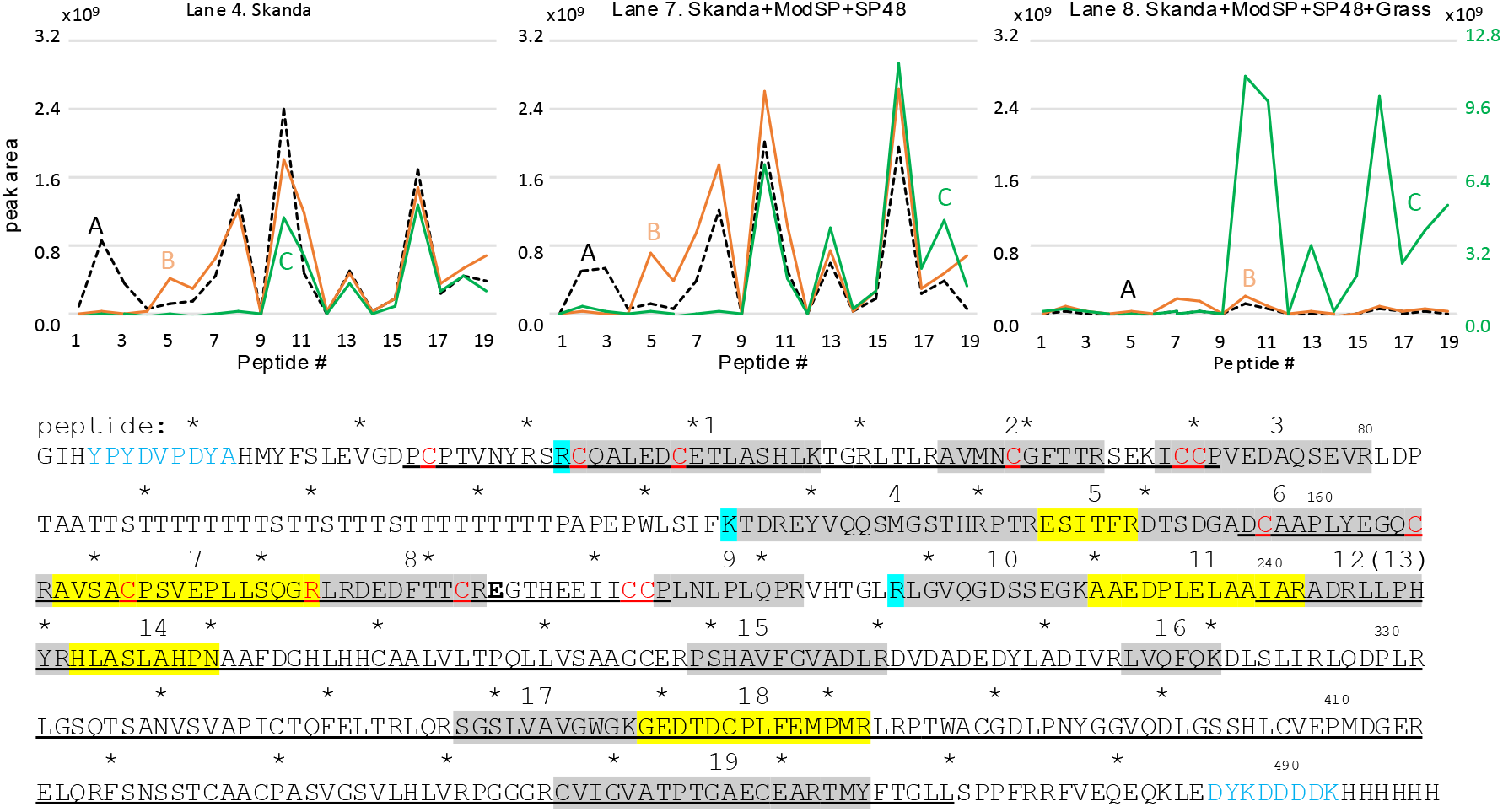
Skanda is an unstable SPH with two clip domains. **(A)** In-gel trypsin digestion and LC-MS/MS analysis of the 85 kDa (**A,** *dashed line*), 44 kDa (**B,** *brown line*), and 35-37 kDa (**C,** *green line*) bands from lane 4 (Skanda alone, *left panel*), lane 7 (ModSP, cSP48, and Skanda, *middle panel*), and lane 8 (ModSP, cSP48, Skanda, and Grass, *right panel*). Locations of peptide-1 through -19 in the recombinant Skanda are shaded *gray* and *yellow.* To aid in residue counting, an asterisk marks every tenth position, and each line ends with a numerical indicator. The residues highlighted in *cyan* indicate that the 85, 44, and 35-37 kDa bands start at or before R32, K125, and R218, respectively. Based on the band sizes and intensities, K125 and R218 may represent the cleavage sites of cSP48, Grass, and a trypsin-like protease released by Sf9 cells. Of note, there is a difference between the sequence for peptide 4 as found in the sequence displayed on Fig. S1: KTDR**D** YV and the sequence of peptide 4 in Table S1: KTDR**E**YV du to the presence of a potential SNP. The cDNA of Skanda was generated by PCR from the BAC03K10. **See Supplementary Document S1.**

**Supplementary Figure 2.**
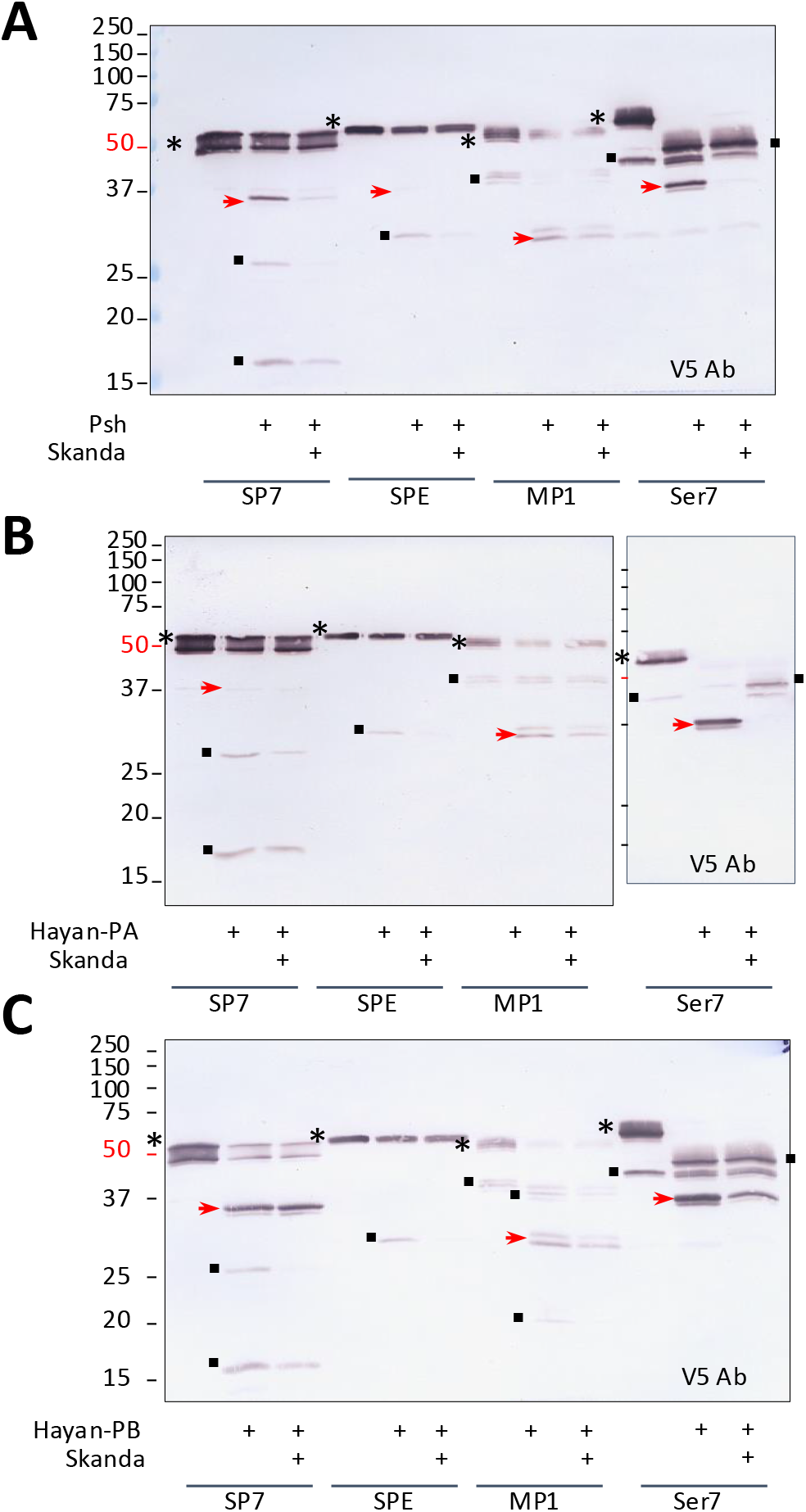
Effect of the Skanda on cleavage activation of *Drosophila* SP7, SPE, MP1, and Ser7 precursors by Psh. Effect of the Skanda on cleavage activation of *Drosophila* SP7, SPE, MP1, and Ser7 precursors by Psh (**A**), Hayan-PA (**B**), or Hayan-PB (**C**). (**A**) ModSP (0.5 μg), proSP48 (0.1 μg), proGrass (0.1 μg), Skanda (0.8 μg), proPsh (0.1 μg), and precursor of SP7 (0.4 μg), SPE (0.5 μg), MP1 (0.2 μg), or Ser7 (0.2 μg) were incubated in 25 μl buffer A at 37 °C for 1 h. The reactions and controls were subjected to 12% reducing SDS-PAGE and immunoblot analysis using 1:1000 diluted antibody against the V5 tag, located at C-terminus of SP7/SPE/MP1/Ser7. In (**B**) and (**C**), proPsh (0.1 μg) was substituted by proHayan-PA (0.4 μg) and proHayan-PB (0.1 μg), respectively. A total of four independent blots were obtained from each repeated experiment for scanning and densitometric measurement. The reductions in SP7 activation (mean ± SEM) caused by Skanda were 56±14% (Psh), 49±19% (Hayan-PA), and 65±18% (Hayan-PB). Reductions in SPE activation were 38±12% (Psh), 43±5% (Hayan-PA), and 23±4% (Hayan-PB). Reductions of MP1 activation were 58±8% (Psh), 44±3% (Hayan-PA), and 90±30% (Hayan-PB). Reductions of Ser7 activation were 9±4% (Psh), 35±1% (Hayan-PA), and 27±13% (Hayan-PB). The precursors (*), catalytic domains (*arrowheads*) and other cleavage products (▪) are indicated, with the catalytic domain *highlighted red*.

**Supplementary Figure 3.**
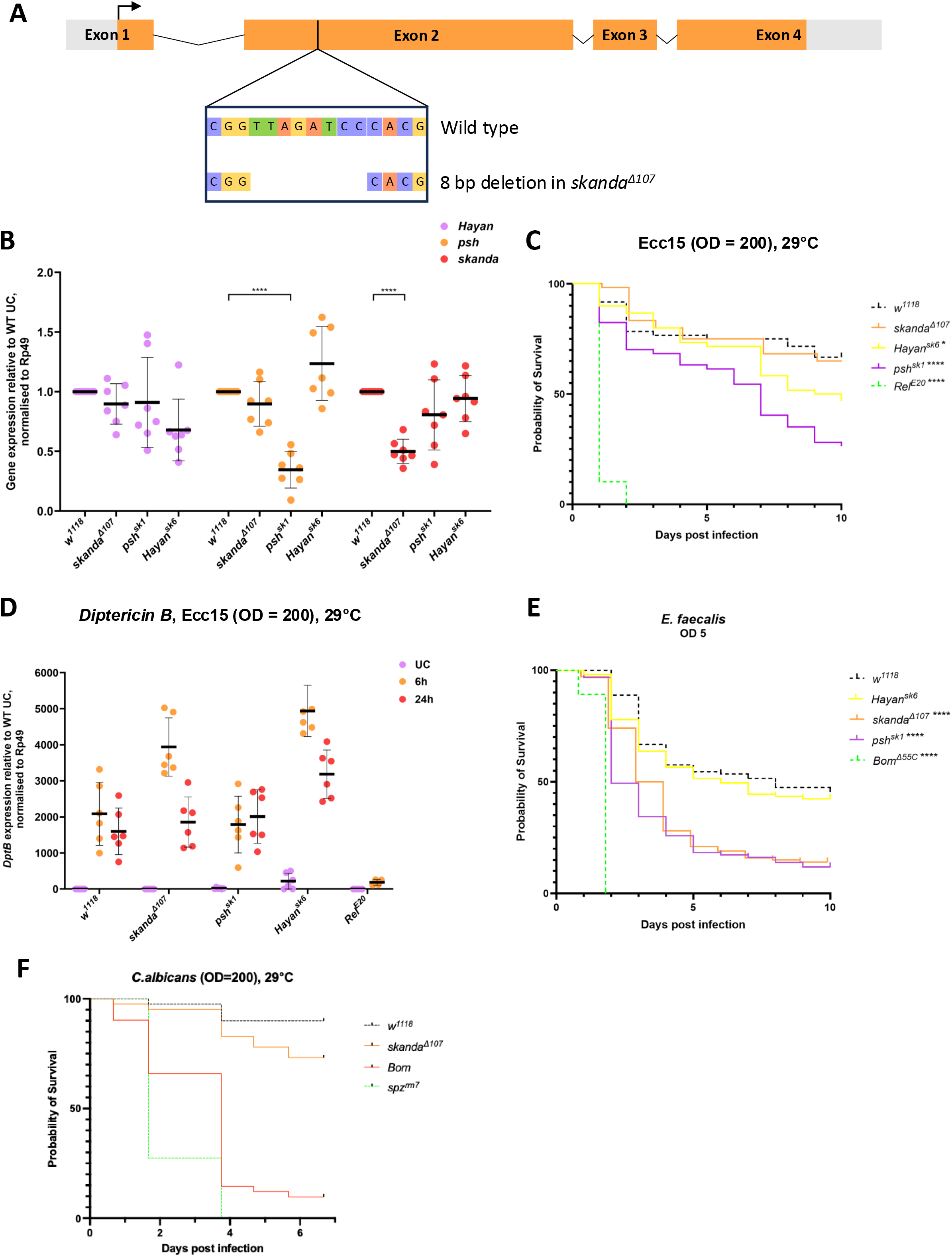
Skanda is not involved in Imd signaling. (**A**) *skanda* is made up of four exons. The *skanda^Δ107^* mutants used in this study contain an 8 bp deletion made by CRISPR-Cas9 in the second exon, disrupting the coding sequence and causing a frameshift. In orange: translated transcript, grey: untranslated region, purple: cytosine, yellow: guanine, green: thymine, red: adenine. (**B**) *Hayan*, *psh* and *skanda* expression levels were observed in the single *skanda^Δ107^*mutant lines. Only expression of the mutated genes was perturbed, and no cis effects were observed. The lower level of *skanda* transcript in the *skanda^Δ107^* flies may be due to RNA decay as the *Δ107* deficiency create a frameshift but does not delete the start of the transcript. (**C**) Survival of s*kanda* mutants upon septic injury with *Ecc15* (O.D. =200). *Rel^E20^* flies were used as a control. All mutants but *Rel^E20^* flies showed wild-type susceptibility to *Ecc1*5. The sudden increase in susceptibility of *Hayan^sk6^* and *psh^sk1^*flies is due to their susceptibility to clean injury at 29°C. (**D**) *DptB* expression of *skanda* mutants was observed at 6 and 24 hours upon pricking with Ecc15 OD600 200. *Rel^E20^*flies were used as a control. *skanda^Δ107^* and *Hayan^sk6^*flies showed slightly higher *DptB* induction at 6 hours post infection, which returned to wild-type levels by 24 hours. *psh^sk1^* flies showed wild-type *DptB* induction. *Rel^E20^* flies showed strongly reduced induction at 6 hours, and were all dead 24 hours post infection. (**E**) Survival of s*kanda* mutants upon septic injury with *E. faecalis* (O.D. =5) at 25°C. *Bom* ^Δ*55*^ flies were used as a positive susceptible control. *Hayan^sk6^* flies showed wild-type survival to *E. faecalis while psh^sk1^and skanda^Δ107^*flies show an intermediate susceptibility. This graph is the pool data of three biological replicates of 20 flies each. (**F**) Survival of s*kanda* mutants upon septic injury with *C. albicans* (O.D. =200) at 29°C. *Bom* ^Δ*55*^ flies and *spz^rm7^and Grass^Herrade^* flies were used as a positive susceptible control. *skanda^Δ107^*flies show an intermediate susceptibility. This graph is the pool data of two biological replicates of 20 flies each. ns = not significant (not shown for Sup. 3B, D). *p < 0.05, **p < 0.01, ***p < 0.001, and ****p < 0.0001.

**Supplementary Figure 4.**
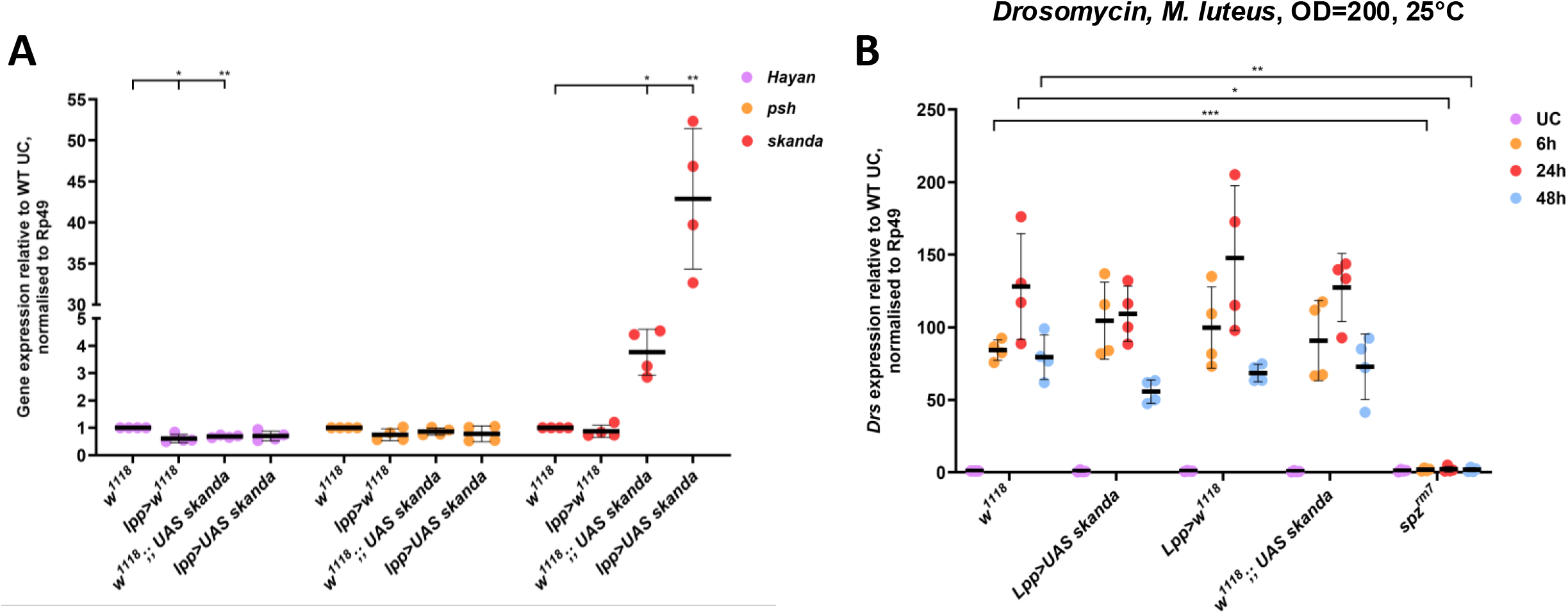
Overexpression of *skanda* does not impact the Toll pathway. (**A**) Driving the expression of *UAS-skanda* with the *Lpp-GAL4* fat body driver leads to a significant overexpression of the gene without impacting the expression of *Hayan* and *psh*. The insertion of *UAS-skanda* also leads to a significant overexpression of *skanda in absence of Gal4*. (**B**) *Drs* expression of flies overexpressing *skanda* was observed 6, 24 and 48 hours after pricking with *M. luteus* OD600 200. *spz^rm7^* flies were used as a control. Flies overexpressing *skanda* had a wild-type Toll response. *Drs* expression was strongly reduced in *spz^rm7^* flies. ns = not significant (not shown for Sup. 4A, B), *p < 0.05, **p < 0.01, ***p < 0.001, and ****p < 0.0001.

**Supplementary Figure 5.**
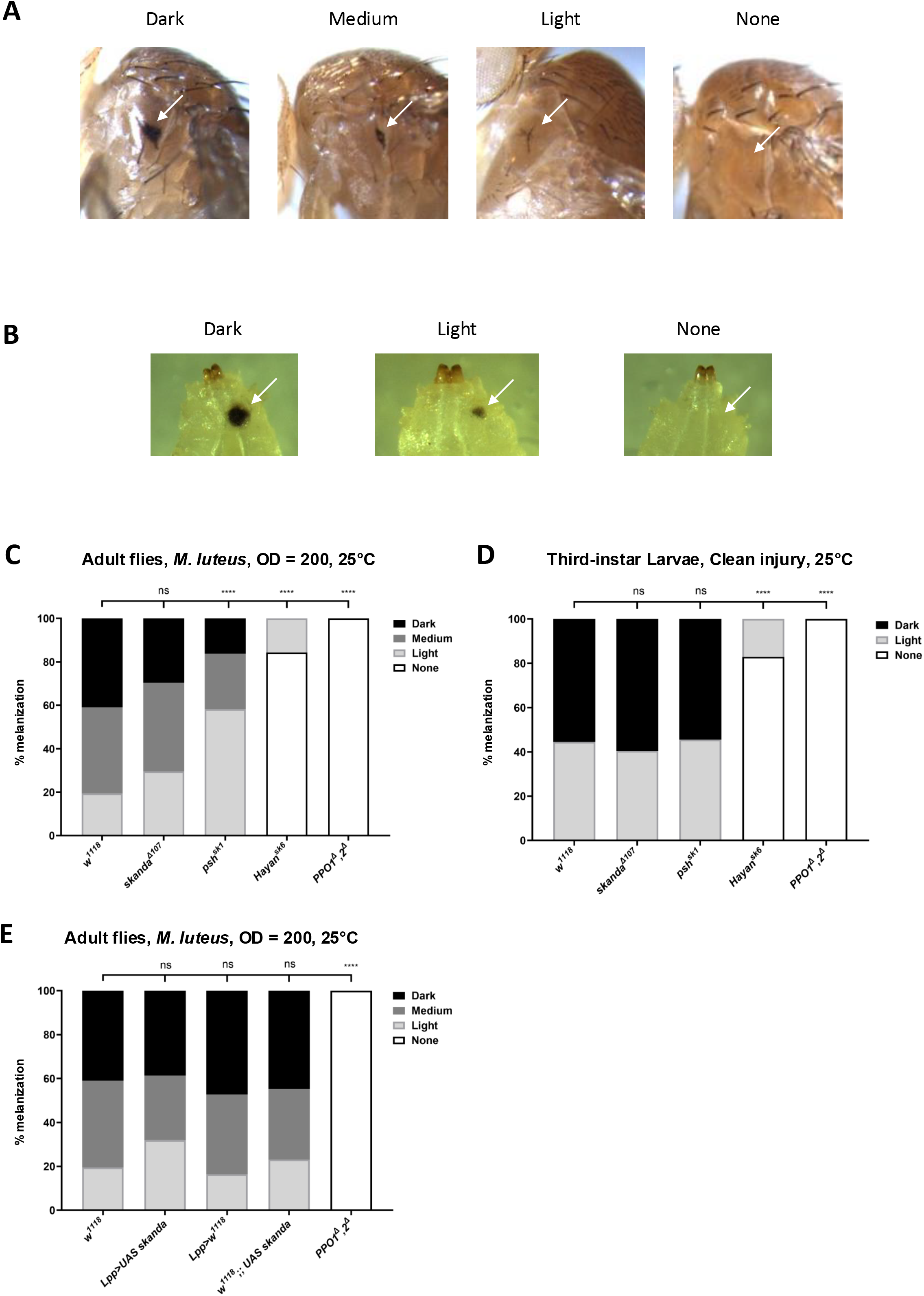
Overexpression of *skanda* impacts cuticular melanization upon clean injury but not septic injury. **(A)** Flies were pricked with a 0.2 mm needle, and cuticular melanization was looked at 16 hours post injury. Intensities were classified as dark, medium, light or no melanization based on the size and color of the scab. White arrow: site of injury and melanization scab. **(B)** Third-instar larvae were pricked with a 0.1mm needle, and melanization was looked at 2 hours post injury. Intensities were classified as dark, light or no melanization based on the size and color of the scab. White arrow: site of injury and melanization spot. Cuticular melanization of flies overexpressing *skanda* was observed 16 hours post **(C)** Cuticular melanization of *skanda* mutants was observed 16 hours post septic injury with a needle dipped in *M. luteus* OD600 200. Dark, medium and light refers to the size and colour of the scab. *PPO1^Δ^,2^Δ^* flies were used as a positive control. *skanda^Δ107^* flies showed wild type melanization, while *psh^sk1^* flies showed reduced melanization. *Hayan^sk6^* flies showed no response to clean injury, but recovered some melanization upon septic injury. As expected, *PPO1^Δ^,2^Δ^* flies showed no melanization. **(D)** Melanization of *skanda* mutant third-instar larvae was observed 2 hours post clean injury with a sterile needle. *PPO1^Δ^,2^Δ^* larvae were used as a positive control. *skanda^Δ107^* and *psh^sk1^* larvae showed wild-type melanization. *Hayan^sk6^*and *PPO1^Δ^,2^Δ^* larvae showed no melanization. **(E)** Cuticular melanization of flies over-expressing *skanda* with the *Lpp-Gal4* driver was observed 16 hours post septic injury with a needle dipped in *M. luteus* OD600 200. Dark, medium and light refers to the size and color of the scab. As expected, *PPO1^Δ^,2^Δ^* flies showed no melanization. ns = not significant, *p < 0.05, **p < 0.01, ***p < 0.001, and ****p < 0.0001.

**Supplementary Figure 6.**
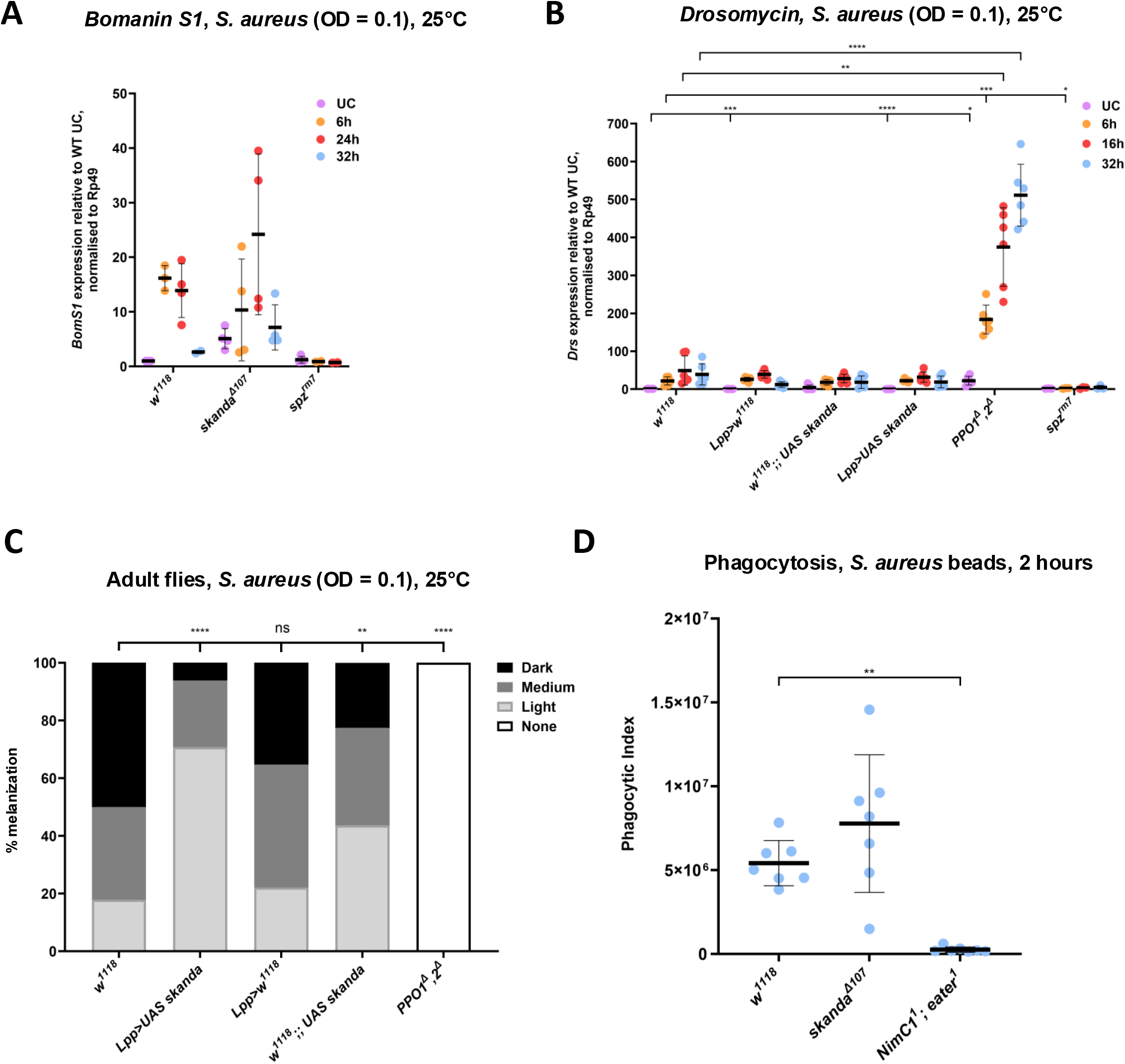
Investigating the mode of host defense conferred by Skanda to *S. aureus*. **(A)** *BomS1* expression in *skanda* mutants was observed 6, 24 and 32 hours after pricking with *S. aureus* at 25°C. *spz^rm7^* flies were used as a positive control. The gene *BomS1* remained inducible in *skanda^Δ107^*flies. **(B)** Cuticular melanization of flies overexpressing *skanda* was observed 16 hours septic injury with a needle dipped in *S. aureus* OD600 0.1. Dark, medium and light refers to the size and colour of the scab. *PPO1^Δ^,2^Δ^* flies were used as a positive control. Flies overexpressing *skanda* showed reduced melanization. *PPO1^Δ^,2^Δ^*flies showed no melanization. **(C)** *Drs* expression of flies overexpressing *skanda* was observed 6, 16 and 32 hours after pricking with *S. aureus* OD_600_ 0.1. *PPO1^Δ^,2^Δ^* and *spz^rm7^* flies were used as controls. Flies overexpressing *skanda* showed wild-type *Drs* expression. *Drs* expression was strongly increased in *PPO1^Δ^,2^Δ^* flies and strongly reduced in *spz^rm7^*flies. **(D)** Phagocytic index of the hemocytes of *skanda* mutant larvae was observed upon incubation with *S. aureus* beads for 2 hours. Flies with mutations in *NimC1* and *Eater* were used a control. Hemocytes of *skanda^Δ107^* mutant larvae were able to phagocytose *S. aureus* beads as well as wild-type hemocytes. ns = not significant (not shown for Sup. 6 A, B). *p < 0.05, **p < 0.01, ***p < 0.001, and ****p < 0.0001.

**Supplementary Figure 7.**
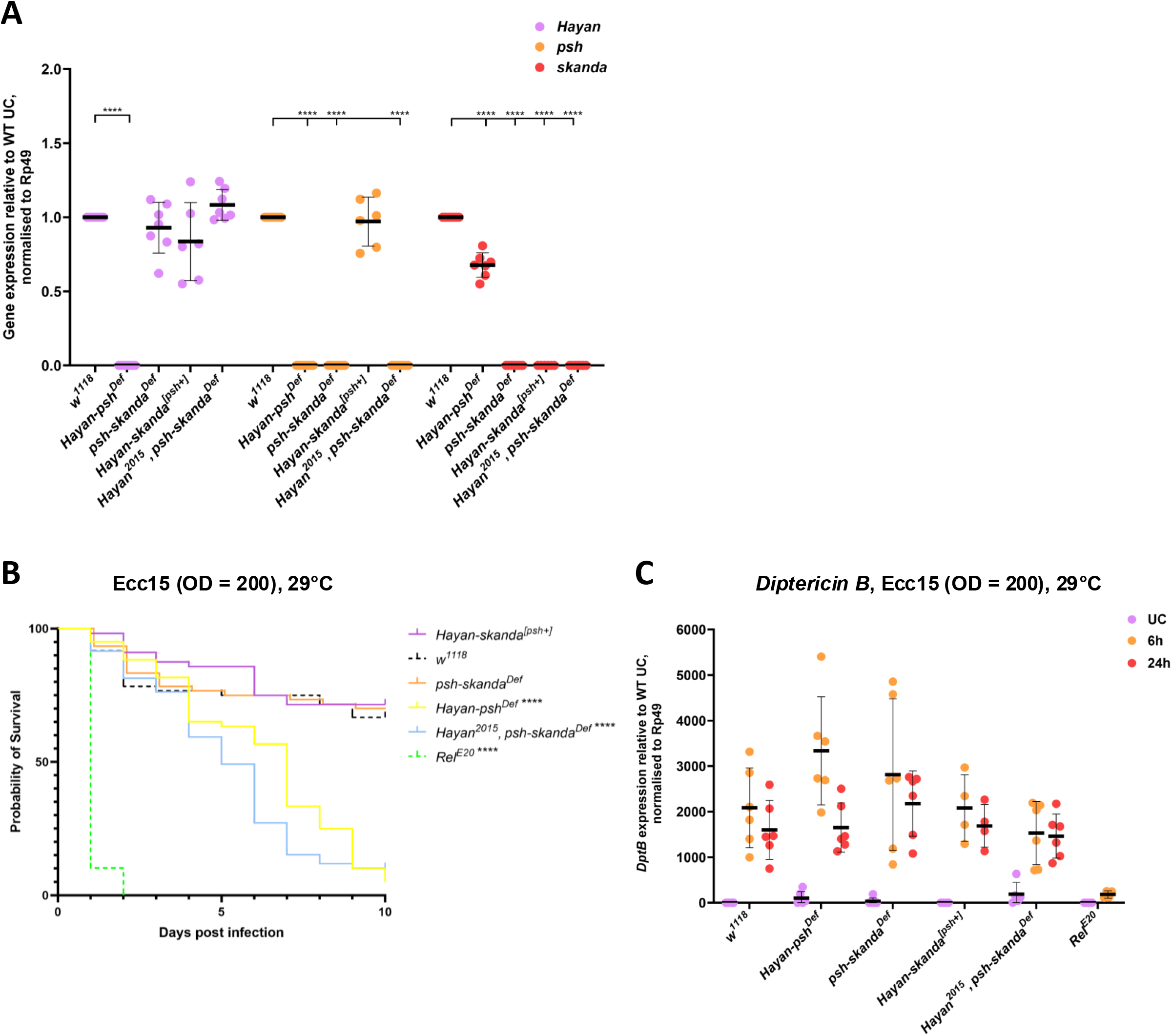
The *Hayan-psh-skanda* gene cluster does not regulate the Imd pathway. (**A**) Levels of expression of *Hayan*, *psh* and *skanda* in the compound mutant lines normalized on the wild-type. Only expression of the deleted genes was perturbed, and no cis effects were observed. Although non-sense mediated RNA decay is often observed, the expression of Hayan in *Hayan-skanda*^[*psh*+]^ and *Hayan^2015^, psh-skanda^Def^* was expected as *Hayan* is still present, although being mutated. The genomic rescue of *psh* provides a wild-type level of expression of psh in *Hayan-skanda*^[*psh*+]^ flies despite the deletion of the endogenous *psh* gene. (**B**) Survival of *Hayan-psh-*s*kanda* compound mutants was observed after pricking with *Ecc15* OD_600_ 200. *Rel^E20^* flies were used as a control. All mutants showed wild-type susceptibility to *Ecc15*. The marked susceptibility of *Hayan-psh^Def^* and *Hayan^2015^, psh-skanda^Def^* flies is due to their susceptibility to clean injury at 29°C. (**C**) The level of *DptB* expression of compound mutants of the *skanda-psh-Hayan gene* cluster was monitored at 6 and 24 hours upon pricking with *Ecc15* OD_600_ 200. *Rel^E20^* flies were used as a control. All mutants showed wild-type *DptB* induction. *Rel^E20^* flies showed strongly reduced induction at 6 hours, and were all dead 24 hours post infection. ns = not significant (not shown for Sup. A, C). *p < 0.05, **p < 0.01, ***p < 0.001, and ****p < 0.0001.

## Supplement Documents

**Supplementary document S1: Characterization of cleaved Skanda product**

To further characterize Skanda cleavage events, we excised three gel slices (**A**, B, and C) corresponding to the 85, 44, and 35−38 kDa regions from lanes 4, 7, and 8 in **Figure 3B**. After reduction, alkylation, and in-gel trypsin digestion, we analyzed nine samples by LC-MS/MS, identifying 19 peptides that together covered 43% of the Skanda sequence. Proteins in gel slices A, B, and C from lane 4 likely begin at or before Cys^33^, Thr^126^, and Pro^204^, respectively (**Figure S1**). For instance, peptide-1 (C^33^QALEDCETLASHLK) and peptide-4 (T^126^DRDYVQQSIGSTHRPTR) result from tryptic cleavage at Arg^32^ and Lys^125^, possibly facilitated by cSP48, Grass, or a trypsin-like protease derived from Sf9 cells. Meanwhile, detection of peptide-9 (P^204^LNLPLQPR^212^)—a trypsin cleavage between S-carboxamidomethylated Cys^203^ and Pro^204^, which is atypical—may indicate secondary cleavage of E^194^GTHEEIICCPLNLPLQPR^212^ (not detected, possibly due to poor ionization). This suggests that proteins in gel slice C start at or before Arg^193^, implying cleavage by SP48 or Grass. Comparing lanes 4 and 7 revealed a marked increase in the peptide abundance in gel slice C, especially peptide-10, -13, -16, and -18, after trypsin digestion (**Figure S1**). Changes in slices A and B were less pronounced. In lane 8, peptide-1 through -19 were nearly undetectable in slices A and B, whereas their levels in slice C peaked in slice C (**Table S1**). To distinguish cleavage specificity between trypsin, SP48, and Grass—all of which share similar substrate preferences—we subjected the nine samples to chymotrypsin digestion followed by MS analysis. Unfortunately, this approach did not provide definitive evidence to map the cleavage sites of SP48 and Grass.

**Supplementary document S2. Effect of the Skanda on the cleavage of SP7, SPE, MP1, or Ser7 precursor by Psh or Hayan**

*Drosophila* Persephone, Hayan-PA, and Hayan-PB have been shown to activate SP7, Spӓtzle processing enzyme (SPE), melanization protease-1 (MP1), and Ser7 to varying degrees (Shan et al., 2023). To examine how Skanda’s modulation of these activators affects downstream cleavage events, we assessed the processing of these four target proteins in the presence or absence of Skanda. In these experiments, ModSP, cSP48 and proGrass were always added to activate Psh or Hayan PA or Hayan-PB. As shown in **Figure S2A**, the catalytic domains of SP7, SPE, MP1, and Ser7 migrated at approximately 35, 36, 30, and 37 kDa, respectively. Additional cleavage products were observed, including a 30 kDa C-terminal fragment of SPE that appeared more abundant than the full-length catalytic domain. The presence of Skanda significantly reduced cleavage efficiency in several cases. Similarly, both Hayan-PA and Hayan-PB, like Persephone, cleaved proSP7, proMP1, and proSer7 to generate their corresponding catalytic domains and additional fragments (**Figure S2B and S2C**). Consistent with the negative impact of Skanda on the cleavage of Hayan and Psh (**Figure 4A**), Skanda tend to inhibit downstream SP SP7, MP1, SPE and Ser7.

**Supplementary document S3 Skanda does no impact the Imd pathway**

We explored whether Skanda regulates the Imd pathway. Adult flies were challenged by pricking with a needle dipped in a concentrated solution of the Gram-negative bacterium *Pectobacterium carotovora carotovora strain 15* (*Ecc15*), a challenge known to strongly induce the Imd pathway. In contrast to *Relish* (*Relish^E20^*) flies that lack a functional Imd response (Hedengren et al., 1999), *skanda^Δ107^* flies display wild-type induction of the antibacterial peptide gene *Diptericin B* and a wild-type survival to infection with *Ecc15* (**Supp. Figure 3C and 3D**). Thus, Skanda does not impact the Imd pathway. This result was not surprising as the Imd pathway is not known to be regulated by serine proteases, but by intracellular and membrane bound PGRP receptors (Ferrandon et al., 2007; Westlake et al., 2024). It however reveals that loss of *skanda* does not lead to a general weakness of the fly that could impact its host defense.

**Table S1.**
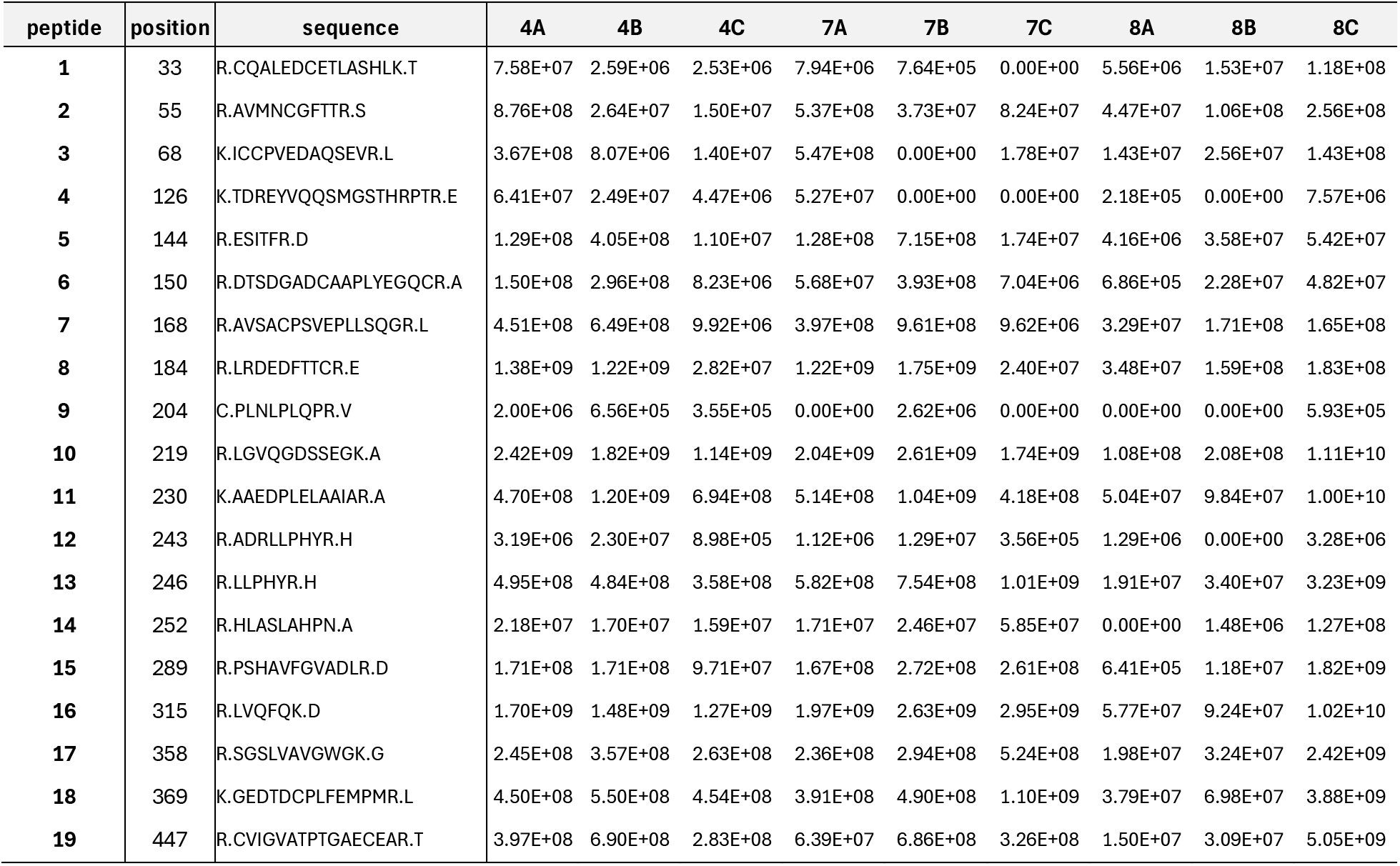
LC-MS/MS determination of the relative abundances of 19 peptides generated by trypsin digestions of Skanda fragments A−C As described in Methods and Materials, lanes 4, 7, and 8 of the SDS-PAGE gel at 85 (A), 44 (B), and 35 kDa positions were excised. The nine gel samples (4A−4C, 7A−7C, and 8A−8C) were treated with TCEP, iodoacetamide, and trypsin. Peptide intensities are listed for deducing the starting sites of bands A, B, and C. Based on the major increases in peak areas (*highlighted yellow*), the 85, 44, and 35-37 kDa bands start at or before R32, K125, and R218, respectively. Based on the band sizes and intensities (Fig. 3), K125 and R218 may represent the cleavage sites of cSP48, Grass, and a trypsin-like protease released by Sf9 cells.

**Table S2.** Structural features of the recombinant proteins and antibodies used to detect them in this study As described in the figure legends, recombinant proteins were detected using a primary antibody against the N- or C-terminal affinity tag. A goat anti-rabbit (GAR) or goat anti-mouse IgG (GAM) conjugated to alkaline phosphatase (AP) was used as the secondary antibody to recognize the antigen–antibody complex and visualize antigens transferred onto the nitrocellulose membrane.

| Name | N-terminal tag | Domain structure | C-terminal tag(s) | 1<sup>st</sup> antibody | 2<sup>nd</sup> antibody |
| --- | --- | --- | --- | --- | --- |
| ModSP | no | LDLa<sub>1-4</sub>-Sushi<sub>1,2</sub>-SP | 6xHis | 6xHis | GAM-AP |
| cSP48 | HA | clip-SP | FLAG-6xHis | FLAG | GAM-AP |
| Grass | FLAG | clip-SP | HA-6xHis | FLAG | GAM-AP |
| Skanda | HA | 2clip-SPH | FLAG-6xHis | 6xHis, FLAG | GAM-AP |
| Psh, Hayan-PA/PB | FLAG | clip-SP | Myc-6xHis | Myc | GAM-AP |
| SPE, Sp7, MP1 | FLAG | clip-SP | V5-6xHis | V5 | GAM-AP |
| Spz | no | Cys knot | 6xHis | Spz C106 | GARat-AP |
| PPO1 | no | I-II-III | no | PPO1 | GAR-AP |

